# Single-cell proteomics reveals proteome remodeling and cellular heterogeneity during NGF-induced PC12 neuronal differentiation

**DOI:** 10.64898/2026.03.25.710659

**Authors:** Arpa Ebrahimi, Shuxin Chi, Jun Yang, Phoebe Y. Lee, Prongbaramee K. Colling, Hailey Mendenhall, Zejing Wang, Chad Giusti, Leonard J. Foster, David A. Hendrix, Claudia S. Maier

## Abstract

Single-cell proteomics (SCP) enables direct measurement of cellular heterogeneity during dynamic biological processes. Here, we applied an SCP workflow to investigate proteome diversity during nerve growth factor (NGF)–induced differentiation of PC12 cells. Differentiated PC12 cells are highly adherent and prone to aggregation, complicating single-cell sample preparation. To address this challenge, sample handling was optimized using gentle dissociation, anti-adhesive conditions, and rapid processing immediately prior to cell isolation. Individual cells were deposited using a refined thermal inkjet (TIJ) dispensing system, enabling accurate single-cell placement with minimal sample loss. Inclusion of the mild nonionic surfactant n-dodecyl-β-D-maltoside (DDM) improved recovery of membrane-associated and other low-solubility proteins. Coupled with high-sensitivity liquid chromatography–ion mobility–mass spectrometry, this workflow consistently quantified approximately 2,000–3,000 proteins per cell across differentiation stages.

Single-cell proteomic profiles acquired over the differentiation time course revealed clear separation between undifferentiated and NGF-treated cells by Day 6. At later stages (Days 4–6), cells further partitioned into two distinct subpopulations with protein expression patterns not evident in bulk measurements. Dimensionality reduction and non-negative matrix factorization identified multiple proteomic states coexisting within the same differentiation stages, characterized by coordinated differences in pathways related to intracellular trafficking, protein translation, and neuronal structural organization. Together, these results show that while global proteome remodeling during PC12 differentiation is captured in both bulk and single-cell data, single-cell proteomics uniquely resolves functionally distinct cellular subpopulations that are masked in population-averaged analyses.

## 2 Introduction

Proteomics provides a direct readout of cellular state by measuring protein abundance and regulation across biological systems. Advances in mass spectrometry–based proteomics have greatly expanded coverage and quantitative performance, enabling detailed characterization of complex cellular processes. However, conventional bulk proteomic measurements report population-averaged signals, which can obscure biologically meaningful variability among individual cells. Single-cell proteomics addresses this limitation by enabling protein-level analysis at cellular resolution, allowing heterogeneity to be directly measured rather than inferred (Kelly, R. T, 2020; Budnik, B., 2018; Vistain, L.F., 2021).

Recent progress in mass spectrometry instrumentation, chromatographic separations, and computational analysis has made it possible to quantify thousands of proteins from individual mammalian cells (Brunner A *et al*. 2022; Specht, H, 2021). These advances have revealed functional heterogeneity that is not always predicted by transcriptomic measurements, highlighting the importance of direct protein-level analysis. Despite this progress, single-cell proteomics remains technically challenging. Low protein abundance, labor-intensive sample preparation, and the need for highly sensitive and reproducible workflows continue to limit throughput and robustness (Kelly R. T., 2020; Momenzadeh, A., 2025). Ongoing efforts to miniaturize and automate sample handling, through nanodroplet processing, microfluidic platforms, and improved acquisition strategies, have begun to address these challenges (Zhu, Y., 2018; Ctortecka, C., 2023; Gebreyesus, S., 2022).

Accurate isolation and handling of individual cells are critical determinants of quantitative performance in single-cell proteomics. Several approaches have been developed, including fluorescence-activated cell sorting (FACS), laser microdissection, and microfluidic or droplet-based platforms (Schoof, E. *et al*., 2021; Zhu, Y. *et al*., 2019; Wang, Y. *et al*., 2024). Each method involves trade-offs between throughput, cell viability, and compatibility with low-volume proteomic workflows. Optimizing cell isolation strategies is particularly important for fragile or highly adherent cell types, such as neurons, where aggregation and cell loss can compromise data quality.

Neuronal differentiation is a tightly regulated process that underlies nervous system development and plasticity. To study proteome remodeling during this process, we used the PC12 cell line, a well-established model of NGF-induced neuronal differentiation. Upon NGF treatment, PC12 cells exit the cell cycle and extend neurite-like processes, adopting features characteristic of sympathetic neurons (Greene, L.A., 1976). This system provides a controlled experimental framework for examining how protein expression programs change as cells transition from a proliferative state to a neuron-like phenotype.

Overall, our optimized single-cell proteomics workflow revealed the phenotype heterogeneity of PC12 cells, resolving transitional states and identifying proteomic signatures that define early, intermediate, and late phases of neuronal differentiation, features that would otherwise be obscured by population averaging.

## 3 Materials and methods

### 3.1 Cell culture

PC12 cells were obtained from ATCC (Part No. CRL-1721; ATCC) and differentiated into neuronal-like cells using 100 ng/mL of Nerve Growth Factor (NGF) (Part No. 86923-98-0; Sigma-Aldrich). Cells were seeded at a density of 2x10^3^ cells/cm^2^ (Hu, R., 2018) in poly-L-lysine (Part No. 0403; ScienCell Research Laboratories) -coated 10 mm culture dishes. After seeding, cells were incubated at 37 °C for 3 hours in RPMI medium (Part No. R8758-100ML; Sigma-Aldrich) supplemented with 5% fetal bovine serum (FBS) (Part No. F4135; Sigma-Aldrich), 10% horse serum (HS) (2 Part No. 6050088; Thermo Fisher Scientific), and 1% penicillin-streptomycin (Part No. 15140122; Thermo Fisher Scientific) to promote adherence to the PLL-coated surface. Following incubation, the medium was replaced with OPTI-MEM (Part No. 15140122; Thermo Fisher Scientific) containing 0.5% FBS to initiate neuronal differentiation. NGF-treated cells were cultured for up to 6 days, with media and NGF replaced every two days to ensure consistent and effective differentiation (Freshney, R. I., 2016). Cells were collected on days 2, 4, and 6 after treatment and dispensed to be used in downstream proteomics analysis.

### 3.2 Single-cell dispensing proteomics workflow

Single cells and reagents were dispensed using the Hewlett-Packard Digital Dispenser (HP-D100), also known as the Uno single-cell dispenser (Tecan, Switzerland) (Figure 1A, Table S1 and S2). Single cells were deposited into 384-well plates (Part No. EP0030129547-25EA; Sigma-Aldrich) pre-filled with 0.5 μL LC/MS-grade water, using C1a cassettes (Part No. 30230841, Tecan) with the medium cell size setting. Then, 1 μL of trypsin digestion solution (Part No. 90057, Thermo Fisher Scientific), containing 1 ng of trypsin/lysC mixture and 0.03% n-dodecyl β-D-maltoside (DDM) (Part No. D4641-500MG; Sigma-Aldrich), was dispensed using D1 cassettes (Part No. 30230843, Tecan) with the surfactant-free setting in the single cell dispenser software (Table S2). Plates were centrifuged at 2,250 × g for 5 minutes to ensure cells and reagents settled at the bottom of each well, reducing sample loss during injection. Digestion was performed by incubating the plate at 37 °C for 2 hours, followed by centrifugation and LC-MS analysis using 384 well plate pre-slit plate sealer (Part No. NC2469150, Fisher Scientific) to prevent contamination and sample evaporation (Figure BS2).

**Figure 1.**
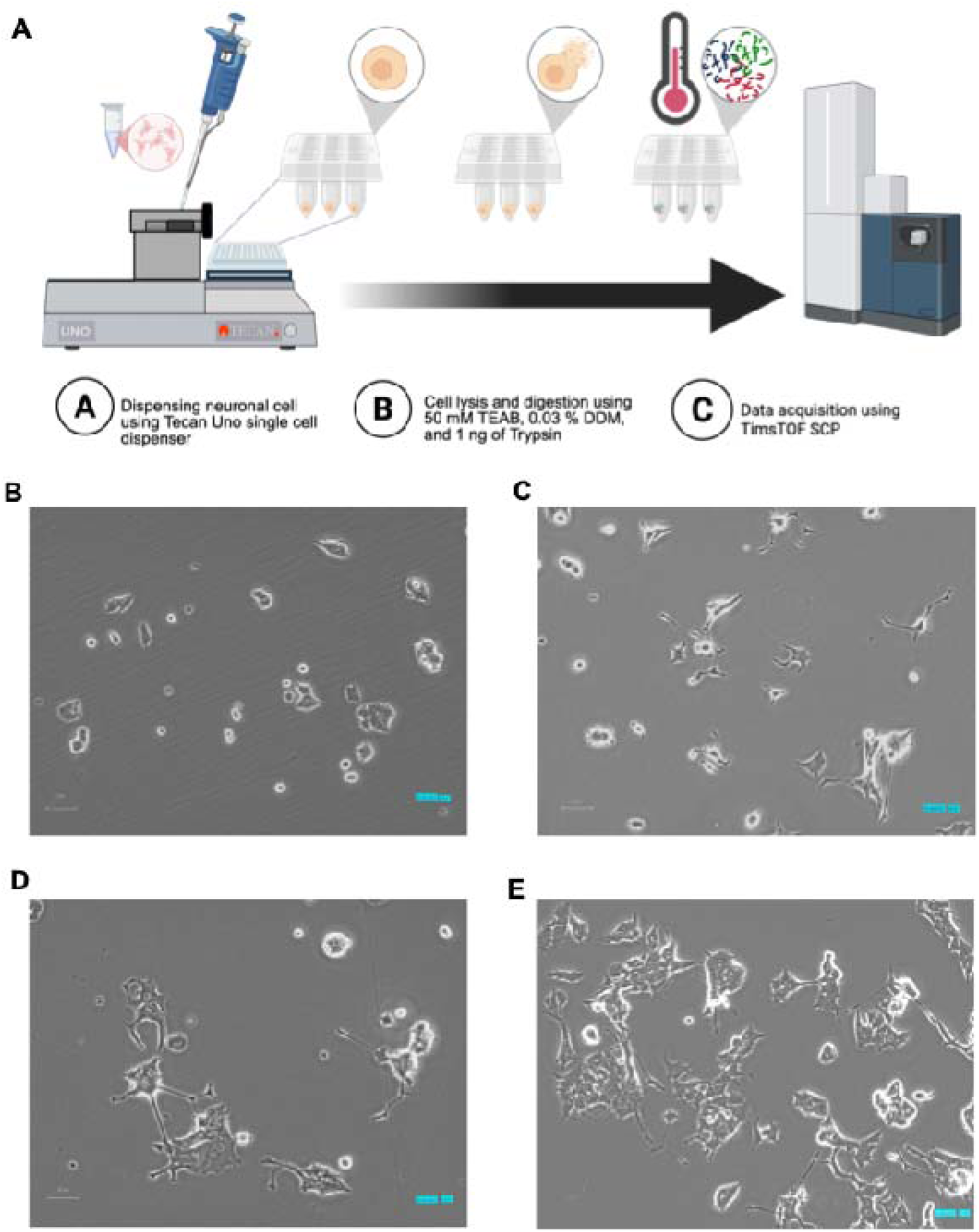
Schematic of single-cell proteomics experimental design and PC12 differentiation. **(A)** Single-cell proteomic sample preparation and proteomics workflow for PC12 cells. The schematic was made with Biorender.com. Representative bright-field microscopy images of PC12 cells across the differentiation time course. **(B)** Undifferentiated PC12 cells at Day 0 without NGF treatment. **(C)** Day 2 following treatment with 100 ng/mL NGF, showing early morphological changes and initiation of neurite extension. **(D)** Day 4 NGF-treated cells exhibiting increased neurite outgrowth and more pronounced neuron-like morphology. **(E)** Day 6 NGF-treated cells displaying extensive neurite networks and mature neuronal morphology. All images were acquired at 20× magnification using bright-field microscopy; scale bar = 50 µm.

### 3.3 Fluorescence imaging to validate single-cell dispensing and neuronal differentiation

Fluorescence imaging was used to assess cell viability and confirm proper single-cell dispensing during neuronal differentiation. Live cells were labeled with Calcein AM (Thermo Fisher Scientific, Part No. 65-0853-39) at an optimized working concentration and incubated for 15 minutes prior to dispensing. Fluorescent signal from viable cells was used to verify cell integrity and handling throughout the procedure. GFP fluorescence images were acquired on the Keyence BZ-X800 microscope using the GFP filter cube (CH2). Images were collected in monochrome mode at high resolution. Because a single field of view did not capture the full well area at the selected magnification, multiple adjacent fields were acquired per well using the multi-point capture function. Autofocus was applied prior to imaging. Individual fields were stitched after acquisition using the Keyence Analyzer software to generate composite images for each well. When enabled, low-photobleach mode was used to restrict excitation light exposure to the time of image capture. Using the Keyence BZ-X800 microscope and fluorescent and bright-field settings, we confirmed the precise isolation of single cells by the single cell dispenser and observed neurite outgrowth indicative of neuronal differentiation. Representative images of stained cells and the labeling mechanism of Calcein AM are shown in Figure S1.

### 3.4 Data acquisition using timsTOF SCP mass spectrometer

Single-cell proteomic analysis was performed using a NanoElute 2 UHPLC system coupled to a Bruker timsTOF SCP operated in dia-PASEF mode (Bruker Daltonics, Germany). Peptides were separated on an Aurora Series Gen3 C18 analytical column (25 cm × 75 µm, 1.7 µm; IonOpticks, Australia) maintained at 50 °C. Mobile phase A consisted of 0.1% formic acid and 0.5% acetonitrile in water, and mobile phase B consisted of 0.1% formic acid and 0.5% water in acetonitrile. Chromatographic separation was performed at a flow rate of 0.3 µL min ¹ using a 15-min gradient.

Mass spectrometric analysis was performed in positive ion mode over an m/z range of 100–1700. The Captive Spray ionization source was operated at 1700 V capillary voltage, 3L/min drying gas, and 200°C drying temperature. During analysis, the timsTOF SCP was operated with Parallel Accumulation-Serial Fragmentation (PASEF) scan mode for DIA acquisition. Trapped ion mobility spectrometry (TIMS) separates ions in the gas phase by balancing an opposing electric field against a constant gas flow, resulting in separation based on ion mobility. In timsTOF instruments, ion mobility is reported as the inverse reduced mobility (1/K, V·s·cm ²), which reflects ion size and gas-phase conformation and serves as a proxy for collision cross section (Michelmann, K *et al*., 2015; Ridgeway, M, E *et al*., 2018). The TIMS analyzer operated in custom mode with an ion mobility range of 1/K = 0.66-1.42, with a 100 ms accumulation time and a 100 ms ramp time. The accumulation time, or trap fill duration, is the time during which ions are stored in the TIMS cell before release; longer accumulation enhances ion collection, which is especially important for sensitivity at single-cell peptide levels. The ramp time indicates how long the instrument takes to scan the full mobility range by lowering the electric field, thereby affecting ion mobility resolution; longer ramps offer higher resolution but slow the duty cycle. These settings resulted in a 9.34 Hz ramp rate. In the dia-PASEF acquisition, a precursor mass range of 400-1000 m/z and a mobility window of 1/K = 0.64-1.37 were used, resulting in a cycle time of 0.96 s. Fragmentation was performed with mobility-dependent collision energies ranging from 20 eV at 1/K = 0.60 to 50 eV at 1/K = 1.60. For each TIMS cycle, 11 dia-PASEF scans were used, each with 3-4 steps. A total of 36 dia-PASEF windows were used, spanning from m/z 299.5 Th to m/z 1200.5 Th, and from ion mobility range (1/K_0_) 0.7 V·s/cm^2^ to 1.3 V·s/cm^2^, with an overlap of m/z 1 Th between two neighboring windows. The collision energy was ramped linearly as a function of mobility value from 20 eV at 1/K_0_ = 0.6 V·s/cm2 to 65 eV at 1/K_0_ = 1.6 V·s/cm^2^.

### 3.5 Single-cell proteomics data processing

Single-cell proteomic data acquired through dia-PASEF were analyzed using DIA-NN (version 1.8.1) (https://github.com/vdemichev/DiaNN) (Demichev, V., 2020) against a Rattus norvegicus FASTA database (UniProt release 2024_01) (UniProt Consortium, 2024) and common contaminants. Bulk libraries were generated using the same method with 10 ng injections. Single-cell samples were analyzed using a spectral library and the Match-Between-Runs (MBR) feature, leveraging bulk samples for alignment. In-silico digestion of a FASTA database was performed using trypsin/P with allowance for one miscleavage. Deep learning-based predictions of MS/MS spectra, retention times (RTs), and ion mobilities (IMs) were enabled to enhance peptide identification. Peptide length ranged between and 30 amino acid residues, precursor charge ranged 2-4, precursor m/z ranged 300-1200, and fragment ion m/z ranged 200-1800. Precursor FDR was set to 1%, with 0 for settings ‘mass accuracy’, ‘MS1 accuracy’, and ‘scan window’. Settings ‘heuristic protein inference’, ‘use isotopologues’, and ‘no shared spectra’ were all enabled. ‘Gene’ was chosen for protein inference parameter along with ‘double-pass mode’ for neural network classifier. Robust LC (high precision) was used for quantification, the RT-dependent mode for cross-run normalization, and smart profiling mode for library generation.

### 3.6 Bioinformatic and statistical data analysis

Single-cell proteomics data were processed using standard filtering, normalization, and statistical procedures. Proteins detected in fewer than 25% of biological replicates were removed prior to downstream analysis. This filtering step was used to reduce the contribution of low-frequency protein identifications. For analyses requiring complete data matrices, missing values were imputed using a k-nearest neighbor (kNN) method based on similarity between samples.

Protein intensity values were normalized on a per-cell basis by dividing each protein intensity by the total protein signal measured within that cell (sum normalization; Figure BS3). Normalized values were log -transformed. Z-score scaling was then applied across proteins. Z-scores were calculated as z = (x_i_ − x) / S, where x_i_ is the protein intensity in a given cell, x is the mean protein intensity across all cells, and S is the standard deviation.

Dimensionality reduction was performed using principal component analysis (PCA) and Uniform Manifold Approximation and Projection (UMAP) (McInnes, L., 2018). Clustering was performed using k-means clustering on normalized protein abundance values. For each cluster, mean normalized protein abundance values were calculated at each differentiation time point.

Non-negative matrix factorization (NMF) was also applied to the filtered protein abundance matrix using non-negative intensity values. Proteins with low detection frequency were removed prior to factorization. Multiple factorization settings were evaluated, and three profiles were retained for downstream analysis. No differentiation time-point labels were included during factorization. Cell-level profile weights were used for visualization, and protein loadings were examined to identify proteins contributing to differences between groups (Lee, D. 1999; Brunet, J., 2004).

Functional enrichment analysis was performed using Metascape (Zhou, Y *et al*., 2019) (v3.5.20250701). Enriched terms with −log (p) ≥ 1.3 were retained for interpretation.

Statistical comparisons of protein abundance between undifferentiated cells and NGF-treated cells at Day 2, Day 4, and Day 6 were performed using unpaired, two-tailed nonparametric tests with α = 0.05. Pairwise comparisons were assessed using the Mann–Whitney U test. Comparisons involving more than two groups were evaluated using the Kruskal–Wallis test followed by Dunn’s post hoc test. Differences between samples prepared with and without n-dodecyl-β-D-maltoside (DDM) were assessed using unpaired, two-tailed Student’s t-tests after confirming assumptions of normality and variance. Precursor-level intensity distributions were compared using log -transformed values and Kruskal–Wallis tests with multiple comparisons correction. All statistical analyses were performed in R using the rstatix package.

## 4 Results

### 4.1 Single-cell proteomics workflow optimization

To establish a robust workflow for single-cell proteomics in this study, several experimental parameters were systematically evaluated using the same analytical platform applied in Chapter 4, including the Tecan Uno single-cell dispenser, NanoElute 2 nanoLC system, and timsTOF SCP mass spectrometer. These optimization experiments were designed to identify conditions that improve protein coverage, reproducibility, and peptide quality while maintaining a workflow that is practical for routine single-cell analysis. Parameters examined included digestion conditions, plate format, sample preparation workflow, chromatographic flow rate, and cell isolation strategy. Data quality was evaluated using multiple complementary metrics, including protein coverage completeness across cells, missed-cleavage rates, peptide length distributions, coefficients of variation (CV) of protein intensities, and peptide-per-protein ratios. The results of these experiments provided guidance for selecting balanced experimental conditions for single-cell proteomic measurements (Chi *et al*., 2026).

Digestion conditions were first evaluated by varying the amount of trypsin used for protein digestion across a range of 0.1–10 ng per well. Increasing the trypsin amount reduced the proportion of missed-cleavage peptides, indicating improved digestion efficiency. However, higher trypsin amounts did not substantially improve overall proteome coverage or completeness across single cells. Instead, intermediate trypsin levels produced the most consistent peptide length distributions and stable protein identifications. These results suggest that moderate digestion conditions provide sufficient enzymatic efficiency while avoiding unnecessary variability in peptide detection. In the optimized workflow, digestion conditions were therefore selected to balance cleavage efficiency with reproducible peptide generation (Chi *et al*., 2026).

Sample handling parameters were also evaluated to assess potential sources of sample loss and technical variability. Comparisons between 96-well and 384-well plate formats showed that low-binding 384-well plates resulted in higher protein completeness and improved reproducibility across cells (Chi *et al*., 2026). The smaller reaction volumes and reduced surface area associated with the 384-well format likely reduce peptide adsorption and sample loss during preparation. In addition, simplified preparation strategies were compared to determine whether additional digestion steps improved performance. A one-step digestion workflow performed comparably to, and in some cases slightly better than, a two-step digestion protocol while reducing processing time and sample handling. These findings support the use of a streamlined sample preparation workflow for single-cell proteomic experiments (Chi *et al*., 2026).

Chromatographic conditions were also examined to determine their impact on data quality. NanoLC flow rates ranging from 100 to 350 nL min ¹ were tested to evaluate the balance between sensitivity and reproducibility. Very low flow rates increased signal intensity for some peptides but were associated with higher variability across single-cell measurements. In contrast, flow rates between approximately 150 and 350 nL min ¹ produced more stable chromatographic performance and lower coefficients of variation. Among the tested conditions, flow rates near 300 nL min ¹ provided a favorable balance between signal intensity, proteome coverage, and quantitative reproducibility. These observations highlight the importance of chromatographic stability in single-cell proteomics experiments, where small variations in peak shape or retention time can influence quantitative measurements (Chi *et al*., 2026).

Finally, different cell isolation strategies were evaluated to determine their suitability for label-free single-cell proteomics. Single-cell dispensing using the Tecan Uno platform was compared with fluorescence-activated cell sorting (FACS). Both approaches produced comparable peptide length distributions and overall protein coverage; however, the Tecan Uno platform provided similar or slightly improved protein completeness while maintaining reproducible quantitative performance. In addition, the Tecan Uno system enables controlled dispensing of individual cells into predefined wells with minimal handling, which simplifies experimental setup and reduces the potential for sample loss during isolation (Chi *et al*., 2026).

Together, these optimization experiments demonstrate that reliable single-cell proteomic measurements can be achieved using a simplified and balanced workflow that minimizes handling steps while maintaining efficient digestion and stable chromatographic separation. Based on these results, the single-cell proteomics experiments described in Chapter 4 were performed using moderate trypsin digestion conditions, a one-step preparation workflow in low-binding 384-well plates, nanoLC flow rates near 300 nL min ¹, and single-cell isolation using the Tecan Uno dispenser. Detailed results of these workflow optimization experiments, including the effects of trypsin amount, plate format, digestion workflow, chromatographic flow rate, and cell isolation strategy, are described in the associated methodological study (Chi *et al*., 2026).

### 4.2 Nanoliter-scale single-cell proteomics workflow for PC12 cells

Building on this, our earlier work established a nano-liter-scaled proteomics workflow using HP’s thermal inject dispenser (Stanishevski, *et al*. 2024). Optimized in HEK-293 cells, this non-contact approach proved robust and broadly applicable to other mammalian cell types, reducing cell damage and contamination and supporting efforts toward accessible, high-throughput single-cell proteomics (Stanishevski, *et al*. 2024; Sanchez-Avila, *et al*., 2023). Using the same single-cell dispenser, we next optimized an already robust workflow for preparing neuronal cells for single-cell proteomics. For high sensitive proteomic analysis, we optimized <u>D</u>ata-Independent <u>A</u>cquisition (DIA) combined with <u>P</u>arallel <u>A</u>ccumulation–<u>S</u>erial <u>F</u>ragmentation (PASEF). This acquisition mode combines advantages of DIA, such as fragmenting all ions within a defined m/z window and providing comprehensive deep coverage, and PASEF, which uses trapped ion mobility spectrometry (TIMS) to accumulate ions in parallel and then fragments them serially based on ion mobility separation.

A single-cell proteomics workflow was applied to PC12 cells treated with NGF and sampled across the differentiation time course (Figure 1A). Over time, cells exhibited morphological changes consistent with neuronal differentiation, including the appearance of neurite extensions and increased cell body size, which were more apparent at Day 4 and Day 6. Within the same time points, individual cells showed variable morphological features, with some cells displaying extensive neurite outgrowth while others retained a more undifferentiated appearance (Figure 1B-E).

### 4.3 Single-cell handling and dispensing optimization for PC12 cells

Processing PC12 cells, particularly after differentiation, was challenging because they are fragile and highly adherent to one another. To preserve cell viability and minimizing aggregation, the differentiated PC12 cells were harvested using Versene, an EDTA-based dissociation solution that chelates Ca² ions to disrupt calcium-dependent cell–cell and cell–substrate adhesion without enzymatic digestion (Takeichi, M., 1991), followed by a 10-minute incubation at 37°C, and then washed with DPBS containing 2 mM EDTA to reduce cell clumping. Before loading into the dispenser, the suspended cells were filtered through a 35 µm cutoff strainer (Part No. 08-771-23, Fisher Scientific) to remove aggregates and debris.

This is a critical step that prevents nozzle clogging and ensures smooth droplet trajectories. Another challenge with differentiated neurons, such as PC12, is their irregular sizes and fragile neurite extensions. To solve this, we systematically tested nozzle settings to ensure precise single-cell deposition without damaging cells (Figure S1, S2). Using the “sticky cell” mode in the Tecan UNO software further improved droplet adhesion, stabilized dispensing of larger or semi-adherent cells, and prevented doublet formation (Table S2). Dispensing accuracy was evaluated with a Keyence microscope (Figure S1). Microscopic inspection confirmed that approximately 85% of wells contained single, viable PC12 cells, demonstrating the high precision and reproducibility of the optimized Tecan UNO settings. These improvements, combining mechanical, software, and biological adjustments, enabled reliable single neuronal cell dispensing, reduced doublet formation, and provided high-quality input material for single-cell proteomic profiling of neuronal differentiation. A total of 163 single PC12 cells were analyzed across four differentiation stages, including Day 0 (N = 35), Day 2 (N = 36), Day 4 (N = 42), and Day 6 (N = 50) (Table S3). To ensure data quality and instrument stability throughout the acquisition, blank injections and 250 pg HeLa digest quality control samples were acquired after every 10 single-cell injections.

### 4.4 Single-cell proteomics dataset overview and quality control

Quality control of the DIA data across all time points demonstrated consistent, reliable instrument performance. Figure 2A-D shows that most proteins were identified by a single peptide, with fewer proteins supported by multiple peptides, which is common in single-cell datasets. The distribution of precursor-level coefficient of variation (CV) across time points (Figure 2E) showed consistent quantitative performance throughout differentiation. All stages, from undifferentiated (Day 0) to fully differentiated cells (Day 6), displayed CV distributions peaking around 30-40%, indicating reliable reproducibility and stable precursor quantification across replicates. The similar shapes and widths of these distributions over time confirm minimal technical variability, while slight broadening at later stages likely reflects increased biological heterogeneity as cells develop neuronal phenotypes. A small percentage of precursors with CV values above 75% correspond to low-abundance features near the detection limit, which is common in single-cell datasets (Tyanova, S. *et al*., 2016).

**Figure 2.**
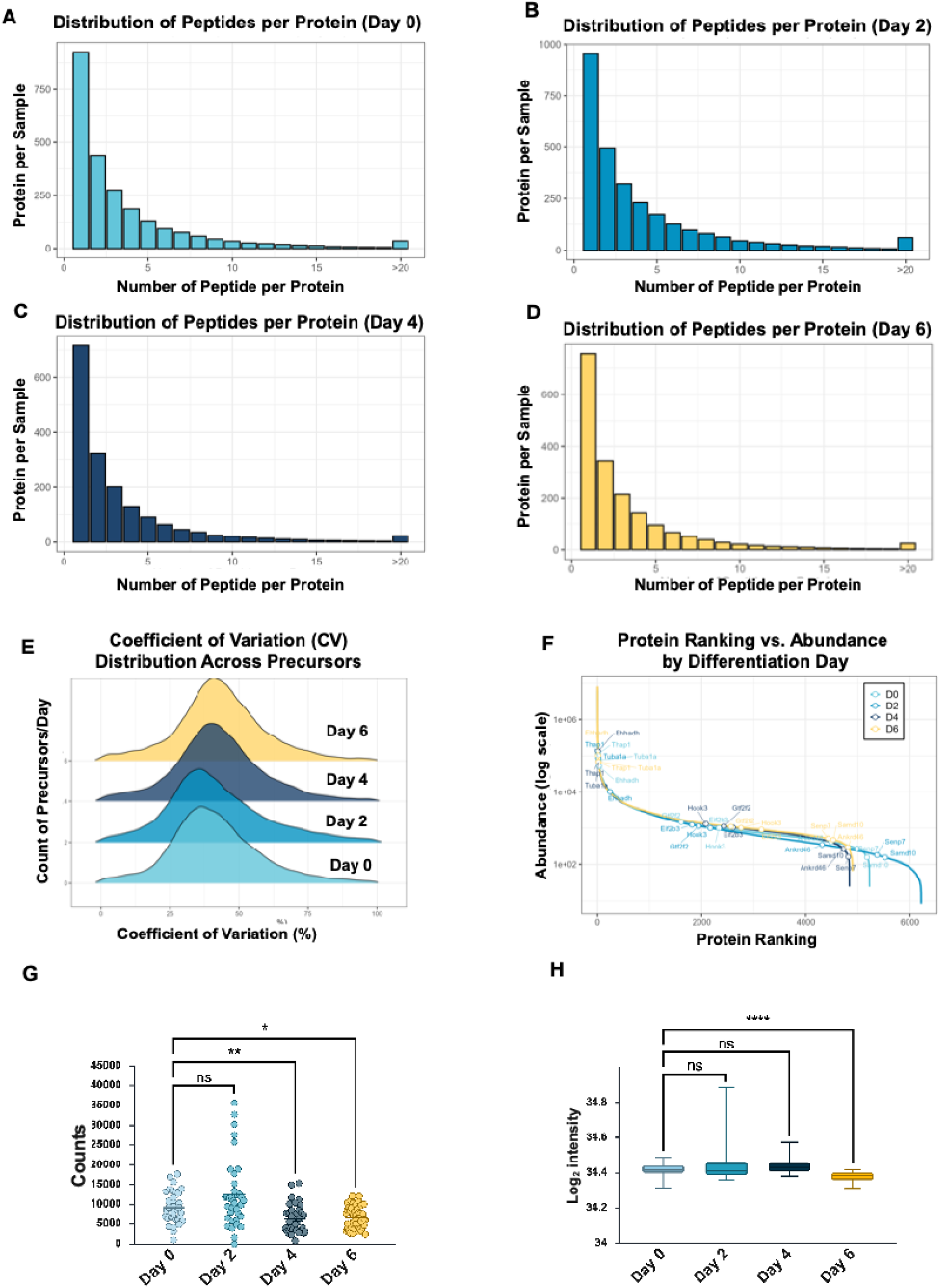
Comprehensive quality control evaluation of single-cell DIA-NN proteomic data. (A–D) Distribution of peptides per protein across Days 0, 2, 4, and 6, showing a right-skewed pattern consistent with single-cell proteomics measurements. **(E)** Distribution of precursor-level coefficients of variation (CVs) across differentiation stages, indicating stable quantitative reproducibility with peak CV values of approximately 30–40%. **(F)** Representative high-, medium, and low-abundance proteins illustrating the broad quantitative dynamic range and consistent detection across differentiation time points. **(G)** Precursor counts per single cell, confirming reproducible identification depth across all differentiation stages, with a small number of Day 2 doublets. **(H)** Log intensity distributions per cell following normalization, demonstrating a uniform quantitative range and minimal run-to-run variation. Statistical analysis (G–H): Group differences were assessed using Kruskal–Wallis tests (nonparametric), followed by Dunn’s multiple comparisons test with adjusted p-values. For panel (H), Kruskal–Wallis testing revealed significant differences across groups (H = 57.68, p < 0.0001). Significance is indicated as ns; *p < 0.05; **p < 0.01; ***p < 0.001; ****p < 0.0001.

To evaluate the quantitative dynamic range of the dataset, representative proteins across high, medium, and low abundance levels were analyzed (Figure 2F). High-abundance proteins, such as Tuba1a, Ehhadh, and Thap1, mainly involved in cytoskeletal structure and energy metabolism, were consistently detected at all time points. Medium-abundance proteins (Eif2b3, Gtf2f2, Hook3), linked to transcriptional and translational functions, and low-abundance signaling or regulatory proteins (Samd10, Ankrd46, Senp7) were also reliably quantified. Overall, the dataset covers a dynamic range of approximately four orders of magnitude, capturing both highly abundant and low-abundance proteins in a single analysis.

The number of precursors identified per cell (Figure 2G) confirms the reproducibility and consistency of DIA acquisitions across the neuronal differentiation timeline. Median precursor counts remained stable from Day 0 to Day 6, indicating uniform detection depth and consistent instrument performance. A small subset of Day 2 cells had unusually high precursor counts, likely due to two-cell dispenses or doublets that increased signal intensity. Differences in precursor counts per single cell across differentiation time points (Day 0, Day 2, Day 4, and Day 6) were assessed using a nonparametric Kruskal–Wallis test because of non-normal data distributions and unequal variances among groups. When a significant overall effect was observed, Dunn’s multiple-comparisons test was applied for post hoc pairwise comparisons with adjusted p-values.

Statistical significance was evaluated at α = 0.05 (Table BS4). Log precursor intensity distributions differed significantly across differentiation stages (Kruskal–Wallis test, H = 57.68, p < 0.0001). Post hoc Dunn’s multiple-comparisons test identified significant differences between Day 6 and all other time points (adjusted p < 0.0001), whereas no significant differences were observed among Day 0, Day 2, and Day 4 (adjusted p > 0.05) (Table S5). The log intensity distributions of precursor abundances across all DIA runs (Figure 2H) were highly overlapping, suggesting consistent signal ranges and minimal variation between runs.

### 4.5 Single-cell proteomics reveals neuronal proteins masked in bulk measurements at late stages of differentiation

Protein ranks from the bulk and single-cell datasets were converted to percentile scores using the formula percentile = 1 − ((rank − 1) / N), where N represents the total number of detected proteins in each dataset. This transformation places both datasets on the same 0–1 scale (1 indicating highest relative abundance), allowing direct comparison despite differences in proteome depth. The percentile values were then matched by protein identifier and visualized in a scatter plot with bulk percentiles on the x-axis and single-cell percentiles on the y-axis. A dashed identity line (y = x) indicates proteins with equivalent relative abundance between modalities. Proteins were ranked according to the difference between bulk and single-cell percentiles to identify those that appear highly abundant in bulk but relatively low in single-cell measurements (Figure S4).

Among the proteins with the largest bulk–single-cell divergence, several are directly relevant to neuronal differentiation and synaptic biology. Cend1 has been shown to promote neuronal lineage progression and differentiation (Baxevanis *et al*., 2010). Baiap2 (IRSp53) is a regulator of actin dynamics and filopodia formation and has established roles in neurite outgrowth and synaptic structure (Chou *et al*., 2017). Pick1 is involved in AMPA receptor trafficking and synaptic plasticity (Xia *et al*., 2000), and Mapk8ip3 (JIP3) functions in axonal transport and neuronal development (Kelkar *et al*., 2000). At the same time, several ribosomal (Rpl29, Rpl37-ps4, Rps7-ps23) and mitochondrial-associated proteins (e.g., Ndufab1, Mrpl3, Ptcd3, Slirp) were also enriched among the bulk-high group, suggesting that structural and bioenergetic components contribute strongly to the averaged bulk signal. Taken together, these results indicate that bulk proteomics amplifies signals driven either by structural programs or by subpopulations of differentiating cells, whereas single-cell proteomics preserves heterogeneity and reveals that many proteins associated with neuronal growth and trafficking are not uniformly abundant across individual cells (Table BS6).

By contrast, single-cell measurements captured these proteins at higher relative abundance, indicating their enrichment within specific subsets of differentiated, neurite-bearing cells. Similar discrepancies between bulk and single-cell proteomic measurements have been reported previously, where proteins linked to defined cellular states are underrepresented in population-averaged data (Kelly, R.T., 2020; Budnik, B. *et al.,* 2018). Together, these observations indicate that bulk measurements underestimate a subset of neuronal proteins that become evident only when cellular heterogeneity is preserved.

### 4.6 Protein identification and protein abundance distribution across differentiation time points

Protein counts were compared across differentiation time points. The number of identified proteins differed between time points (Figure 3A), with fewer proteins detected at Day 4 and Day 6 compared with Day 0 (H = 38.81, p < 0.0001; Table S7). Protein abundance values were visualized by principal component analysis (Figure 3B). Day 0 replicates form a compact cluster. Samples from later time points occupy a larger area of the PCA space. Dispersion is most pronounced at Day 4 and Day 6.

**Figure 3.**
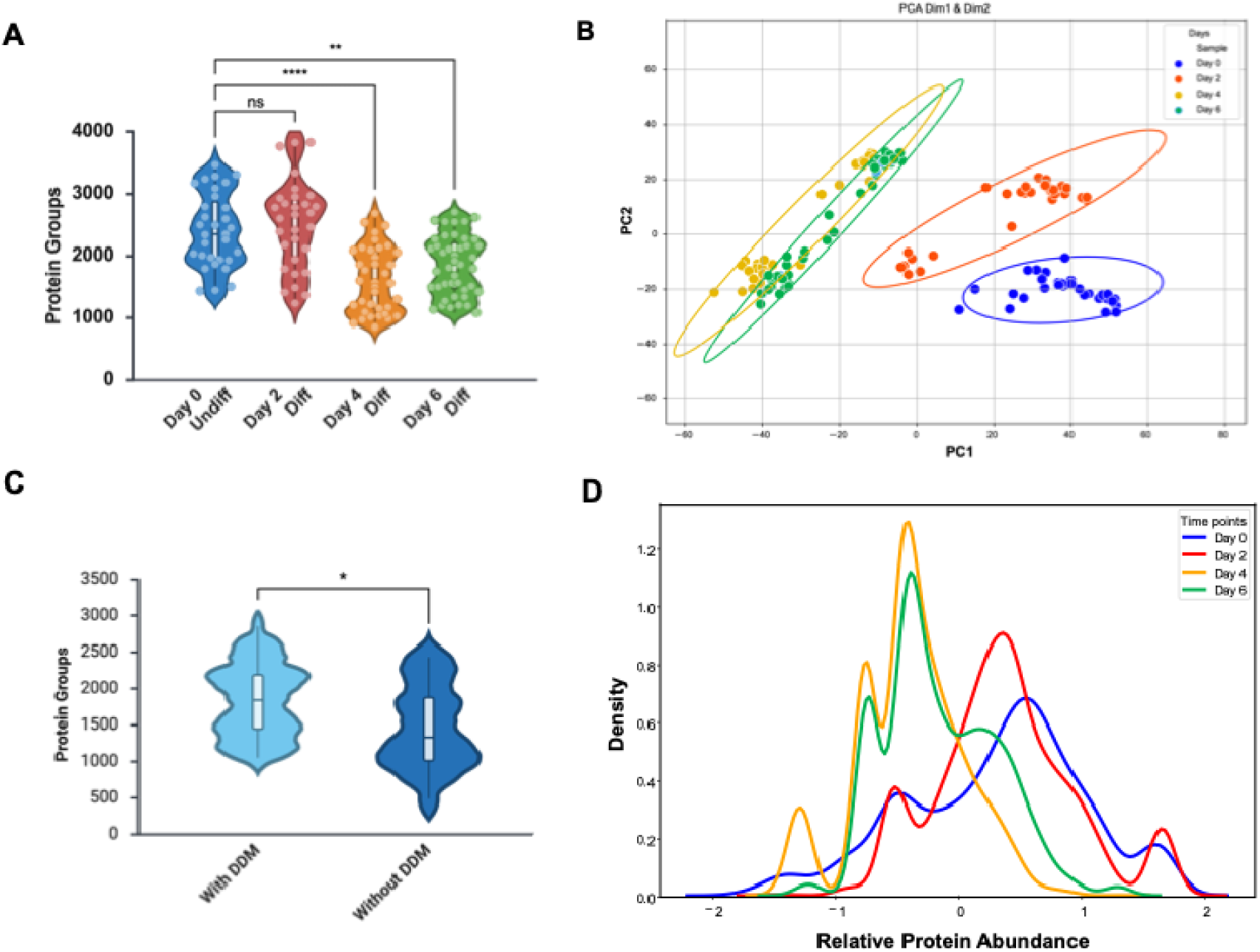
Protein group quantification, Principal Component Analysis (PCA), and density distribution across differentiation time points. **(A)** Protein group identification at different differentiation stages. The number of protein groups identified varies between individual cells at the same time, reflecting cell-to-cell differences in proteome depth during differentiation. Kruskal–Wallis testing was applied due to non-normal distributions. Dunn’s multiple comparisons test was used for post hoc analysis with adjusted p-values. Significance: ns, not significant; **p < 0.01; ***p < 0.0001. (B) Principal component analysis (PCA) of single-cell proteomes collected at Days 0, 2, 4, and 6. **(C)** Violin plots showing the distribution of protein group identifications per single cell for samples prepared with DDM and without DDM. (*t* = 2.61, df = 60, *p* = 0.011) **(D)** Density plot of single-cell proteomic profiles across four differentiation time points. Kernel density estimation summarizes the distribution of proteomic profiles for each time point.

Protein identifications were also compared between samples prepared with and without n-dodecyl-β-D-maltoside (DDM). Samples prepared with DDM show, on average, higher protein counts (Figure 3C). A difference between the two groups was observed by an independent samples t-test (p = 0.011; Table S8). DDM has been reported to improve recovery of membrane-associated and low-solubility proteins in LC–MS workflows (Rabilloud, T., 2009; Chandramouli, K., 2009; Zhang, X. *et al*., 2015; Hughes, C.S. *et al*., 2019). Protein abundance distributions are shown as density plots in Figure 3D. Day 0 and Day 2 show relatively simple distributions. Day 4 and Day 6 show broader distributions with multiple peaks. The distributions at Day 4 and Day 6 are shifted toward lower relative abundance values.

Single-cell protein abundance heatmaps are shown in Figure S5 without clustering (A) and with clustering (B). When ordered by time point, samples show similar overall trends from Day 0 to Day 6. After clustering, subsets of proteins with opposing abundance patterns are visible within Day 2, Day 4, and Day 6 samples. These features are not apparent in the unclustered heatmap.

### 4.7 Effect of DDM on protein recovery in neuronal samples

Comparison of neuronal samples prepared with and without n-dodecyl-β-D-maltoside (DDM) showed clear differences in the proteins recovered (Figure S6A). Proteins enriched in the DDM condition included vesicular and lysosomal proteins such as Cathepsin D and Transferrin, as well as proteins with limited solubility or strong structural associations, including Desmoplakin and Hemoglobin β. Several secreted proteins, including Alpha-fetoprotein and Inter-alpha-trypsin inhibitor heavy chain 4, were also detected at higher abundance when DDM was used (Figure BS6B). Samples prepared without DDM had lower abundances of soluble and membrane proteins, indicating reduced recovery of membrane-proximal and poorly soluble species under detergent-free conditions (Rabilloud, T., 2009; Settembre, C *et al*., 2013; Buszczak, M. *et al*., 2014).

### 4.8 Comparison of Day 0 and Day 2 proteomes

To capture the earliest proteomic changes induced by NGF, we compared protein expression between Day 0 and Day 2, focusing on the top 20 proteins with the largest fold changes and lowest p-values (Figure 4A–C; Table S11). These proteins fall into functional groups that match the pathways highlighted in the enrichment analysis. Lysosomal proteins such as Lamp1 and Ctsl decrease early, matching the “Lysosome” pathway in Figure 4D and reflecting reduced turnover as cells exit a proliferative state (Calegari, F., 2003). Proteins involved in lipid and sphingolipid metabolism, including Asah1 and Msmo1, also decline and align with enriched pathways related to alcohol and modified amino acid biosynthesis, cholesterol metabolism, and other lipid-processing functions (Buszczak, M. *et al*., 2014).

**Figure 4.**
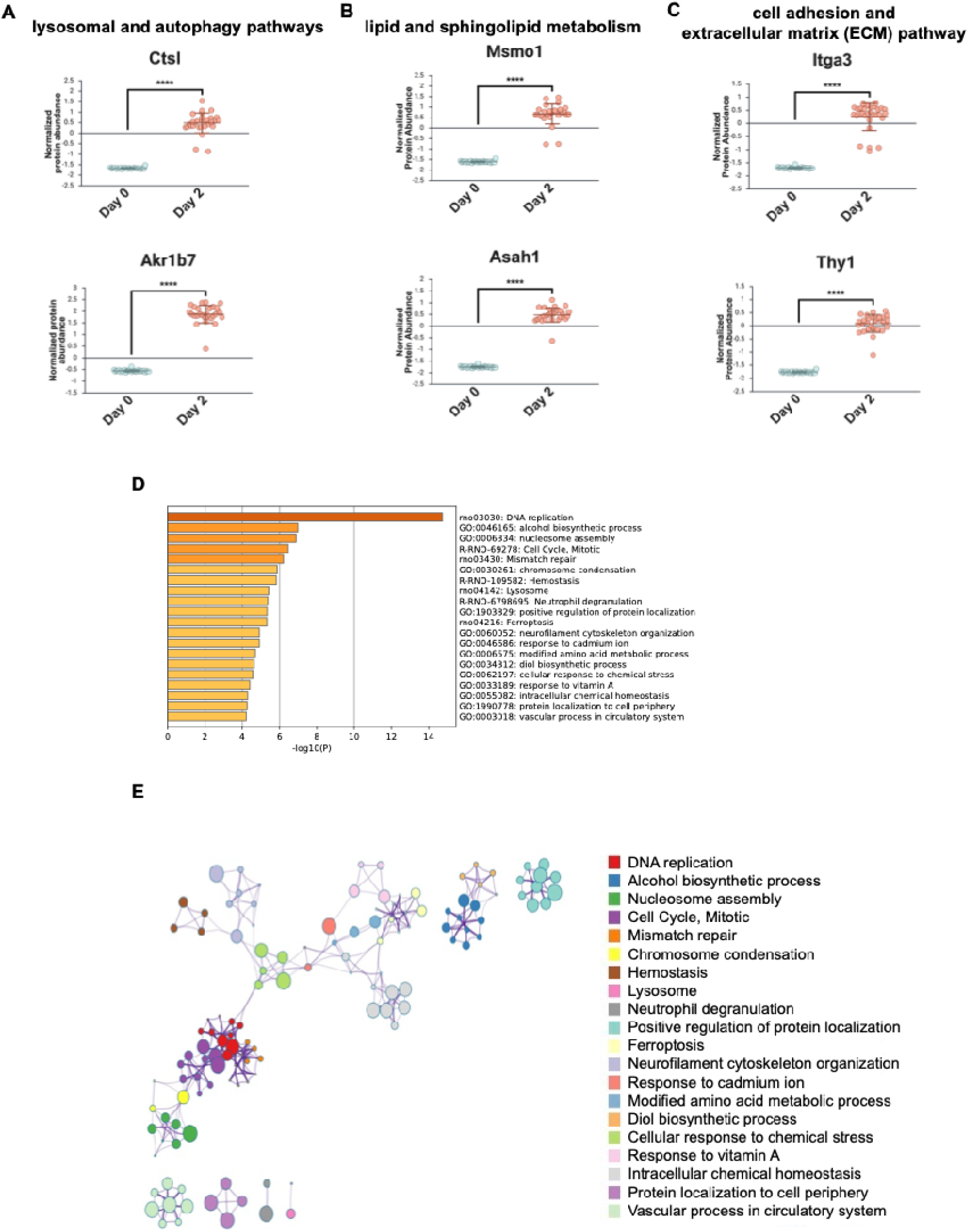
Differential protein expression and pathway enrichment between Day 0 and Day 2. **(A-C)** A representative subset of proteins with the lowest p-values distinguishing Day 0 and Day 2, grouped by functional category (two-tailed Mann–Whitney U test, ****p < 0.001). Horizontal lines indicate group medians. **(D)** Bar plot of enriched biological terms for the input protein lists, colored by adjusted p-value. **(E)** Enrichment network visualization with nodes colored by cluster assignment.

Adhesion-related proteins like Thy1 and Itga3 map to pathways associated with cytoskeletal organization and protein localization to the cell periphery, which fits with the early detachment step before neurite extension (Settembre, C. *et al*., 2013; Ledesma, M.D. *et al*., 2012). Stress- and iron-regulation proteins are enriched in categories such as intracellular chemical homeostasis (Leyton, L *et al*., 2001; Margadant, C *et al*., 2010). The STRING network (Margadant, C., 2010) in Figure 4E groups these pathways into clear clusters, showing how changes in lysosomal activity, lipid metabolism, adhesion, and stress response are interconnected in the early transition toward a neuronal phenotype.

### 4.9 Comparison of Day 0 and Day 4 proteomes

To examine how differentiation advances beyond the early NGF response, we compared Day 0 and Day 4 using the most statistically significant proteins with low p-values and large fold changes (Table BS12) (Figure 5A-B). By this point, most cells have exited the cell cycle, as reflected by significant downregulation of proliferation markers such as Mki67 and Cdk1. These proteins typically peak during the G1–S–G2–M phases, so their decrease indicates that cells are no longer dividing and have fully left the cell-cycle program (Bogdan, A. R. *et al*. 2016). This pattern aligns with the enrichment analysis in Figure 5C, where pathways related to DNA replication, mitotic progression, mismatch repair, and nucleosome assembly are among the most depleted. In contrast, proteins associated with differentiation and cellular remodeling remain elevated at Day 4. Asah1 and Thy1, which were already increased at Day 2, continue to show higher abundance, aligning with pathways involved in membrane organization, lipid metabolism, and protein localization. The rise in Nipsnap3a corresponds with enriched categories related to RNA metabolism, mRNA processing, and intracellular homeostasis, reflecting that transcriptional and metabolic programs are being reorganized as the cells mature (Holt, C. E. *et al*., 2019). The STRING network in Figure 5D groups these enriched terms into distinct clusters, illustrating how shifts in RNA processing, lysosomal function, membrane remodeling, and stress responses fit together to define the Day 4 state. Overall, these patterns show that by Day 4, PC12 cells have largely stopped cycling and are engaging in metabolic, membrane, and RNA-regulatory activities that support progression toward a more defined neuronal phenotype (Hardwick, L. J. A. *et al*, 2015).

**Figure 5.**
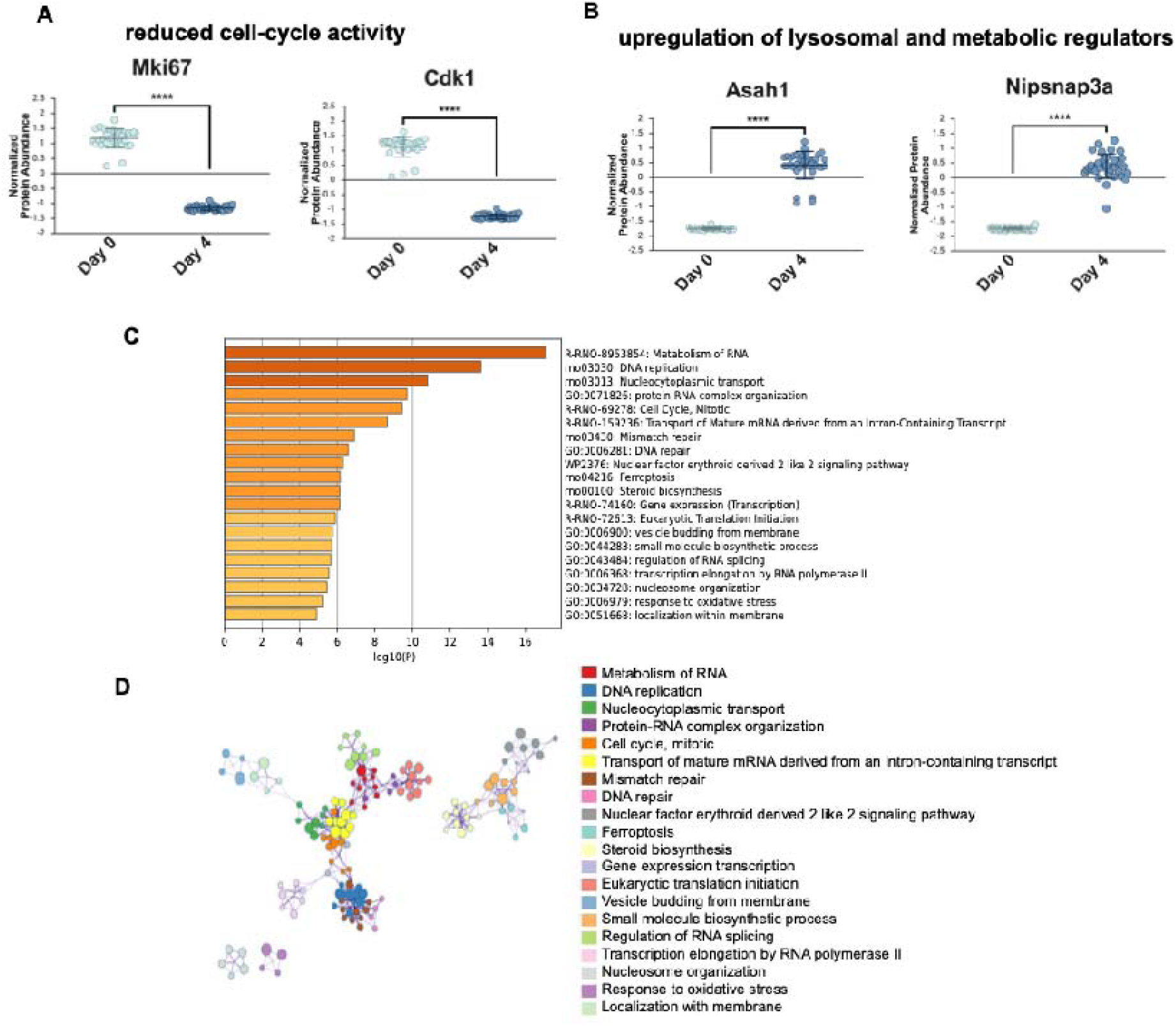
Differential protein expression and pathway enrichment between Day 0 and Day4. **(A**) Single-cell abundances of MKI67 and CDK1 at Day 0 and Day 4. **(B)** Single-cell abundances of ASAH1 and NIPSNAP3A at Day 0 and Day 4. Points represent individual cells; (two-tailed Mann–Whitney U test, ****p < 0.001). Horizontal lines indicate group medians. **(C)** Bar plot of enriched biological terms for the input protein lists, colored by adjusted p-value. **(D**) Enrichment network visualization with nodes colored by cluster assignment.

### 4.10 Comparison of Day 0 and Day 6 proteomes

Comparison of undifferentiated (Day 0) and NGF-differentiated (Day 6) PC12 cells revealed increased levels of protein markers associated with neuronal cytoskeletal organization and maturation. Neuronal intermediate filament proteins NEFM and NEFL, along with INA, Prph, and TUBB2A, showed higher abundances at Day 6 relative to Day 0. These proteins are key components of neurite structure, microtubule dynamics, and neuronal stabilization, indicating substantial cytoskeletal remodeling during differentiation (Yuan, A. *et al*., 2017) (Figure 6A). Pathway enrichment analysis and protein interaction network revealed strong overrepresentation of RNA metabolic processes, mRNA processing, protein folding, and intracellular transport pathways, indicating increased biosynthetic and regulatory activity associated with neurite extension and stabilization (Goncalves, J. T. *et al*, 2016) (Figure 6B-C, Table S13).

**Figure 6.**
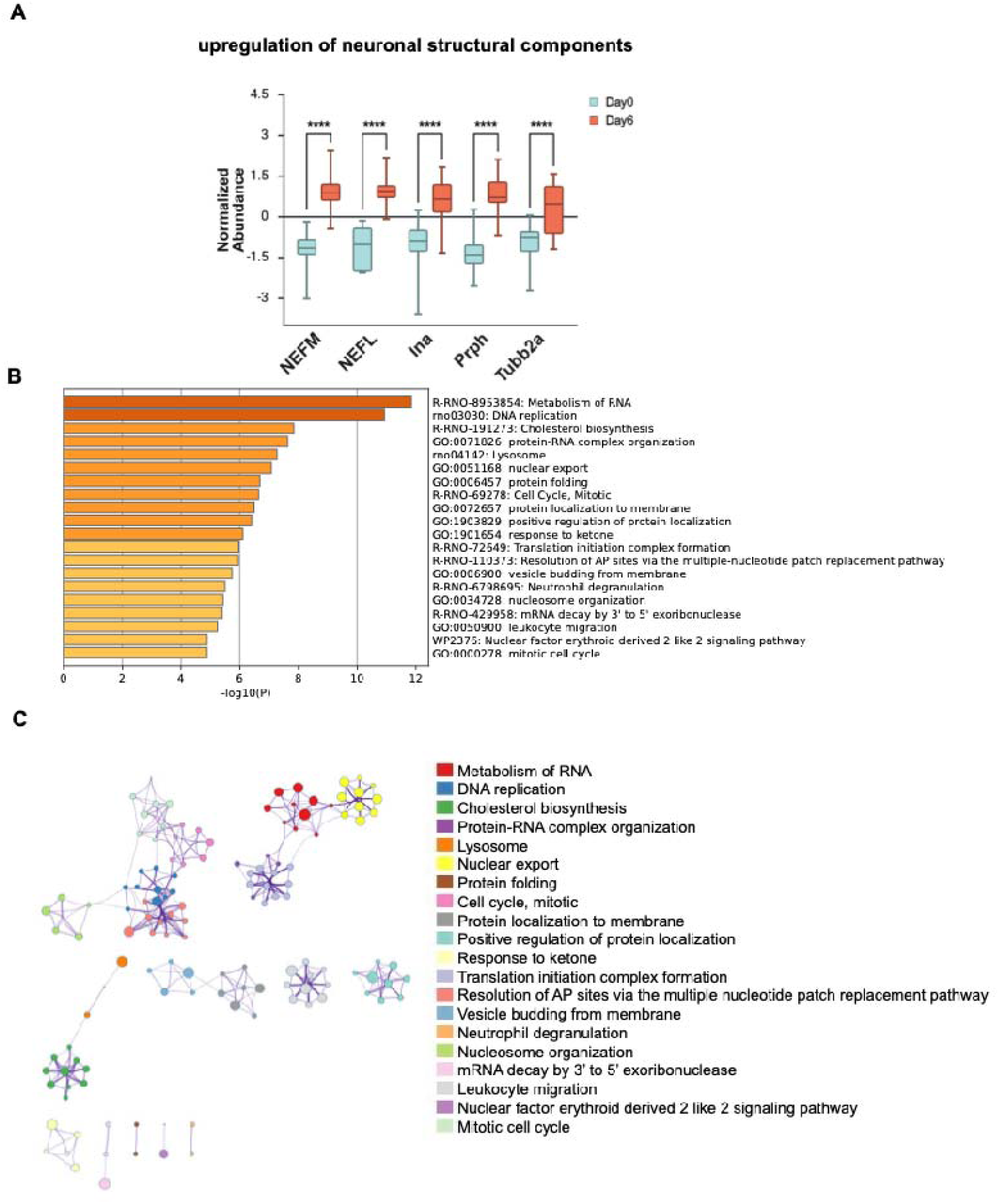
Differential protein expression and pathway enrichment between Day 0 and Day 6. **(A)** Boxplots show the distribution of normalized protein abundances for selected neuronal markers (NEFM, NEFL, INA, Prph, and TUBB2A) at Day 0 (undifferentiated) and Day 6 (differentiated) PC12 cells. (two-tailed Mann–Whitney U test, ****p < 0.001). Horizontal lines indicate group medians. **(B)** Bar graph displaying enriched biological terms across the input gene lists, with bars colored according to adjusted p-value. **(C)** Enrichment network visualization, where nodes are colored based on cluster ID; terms within the same cluster are positioned near each other, indicating shared biological themes.

### 3.11 UMAP projection of single-cell proteomic profiles

To further evaluate neuronal features emerging at later stages of differentiation, we next examined how known maturation markers distribute across the single-cell proteome using UMAP projections (Figure 7). UMAP showed distinct separation between the later time points (Day 4 and Day 6) and earlier ones (Day 0 and Day 2). When we mapped neuronal maturation markers onto this space, proteins such as NEFL, NEFM, TUBA4A, PRPH, and VGF showed higher intensities primarily in the Day 4 and Day 6 clusters (Table S9).

**Figure 7.**
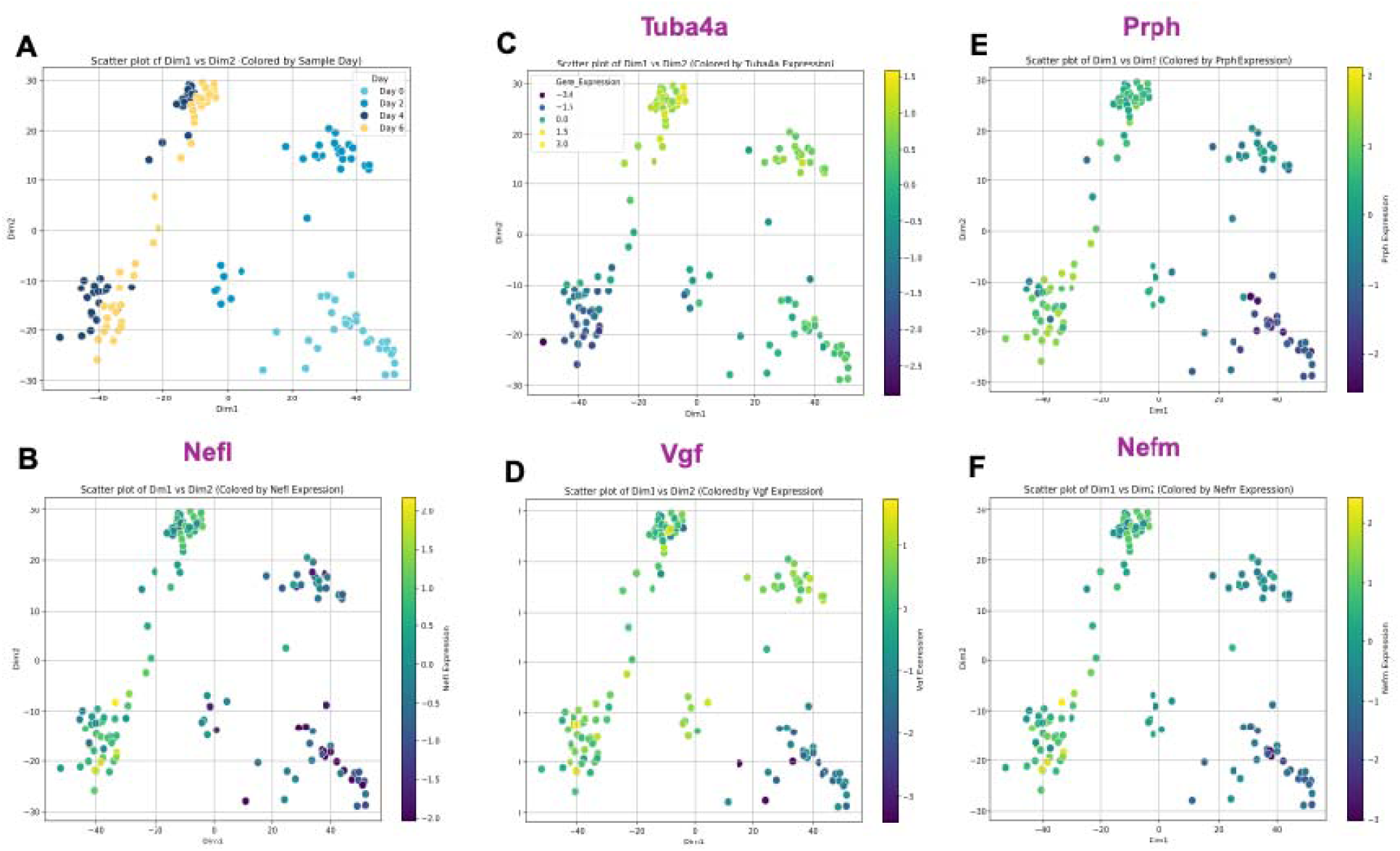
Expression maps of neuronal maturation markers. **(A)** PCA plot of all time-course samples, showing separation of early and late differentiation stages. **(B-F)** Expression patterns of key neuronal maturation proteins: Nefl, Tuba4a, Vgf, Prph, and Nefm. All five proteins show increased abundance in Day 4 and Day 6 clusters.

### 4.12 Temporal protein expression patterns across UMAP-defined clusters

Single-cell protein abundance data from Day 0, Day 2, Day 4, and Day 6 were visualized using UMAP (Figure 8). The resulting embedding separates cells into seven clusters based on similarity in protein abundance profiles. Cells from different time points are present across multiple clusters. For each cluster, relative protein abundance values were summarized by differentiation day. Proteins within clusters show different patterns across time, including decreases, increases, and transient changes between intermediate time points.

**Figure 8.**
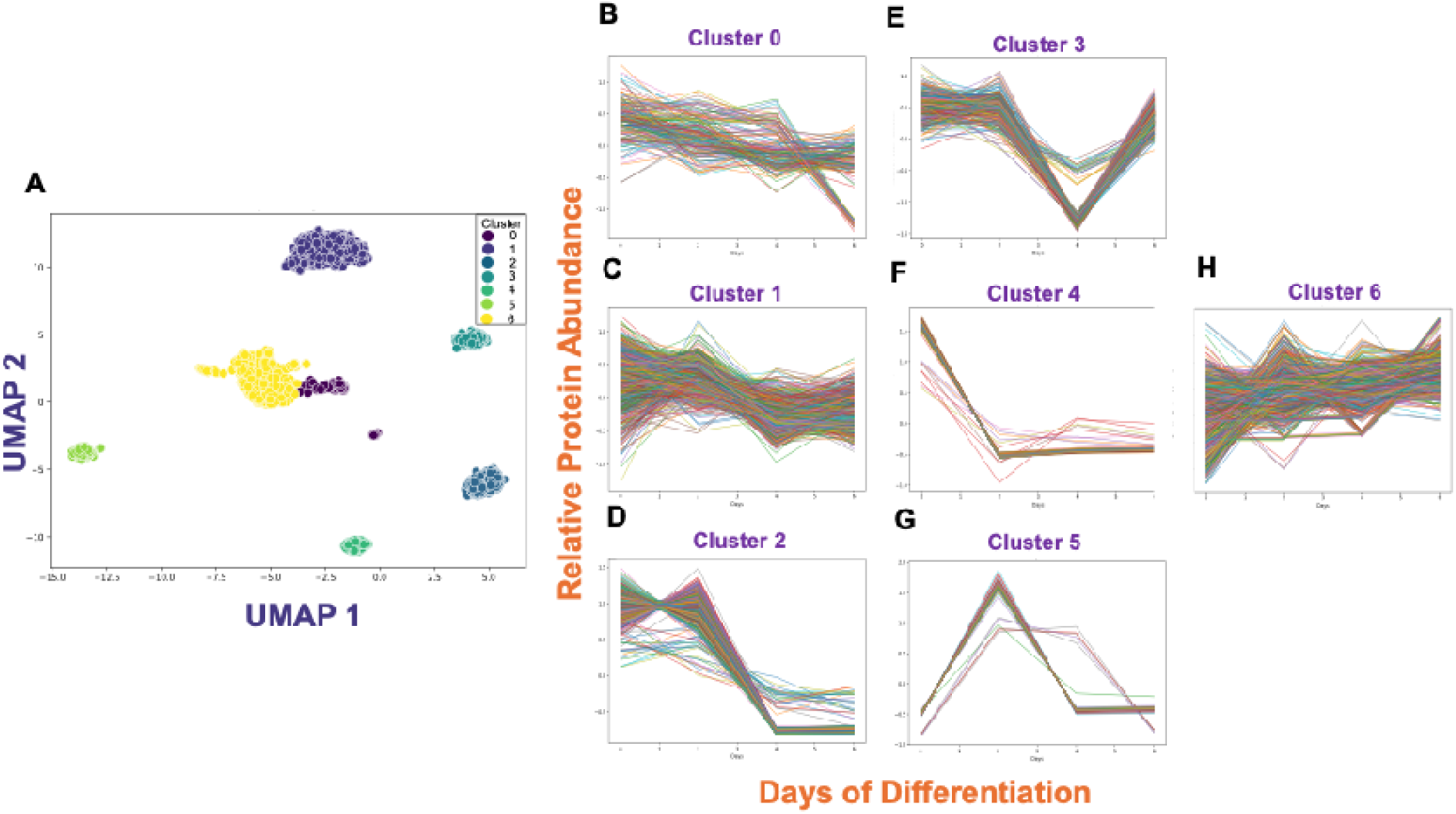
UMAP-based clustering reveals distinct protein expression trajectories during PC12 neuronal differentiation. **(A)** Single-cell proteomic profiles collected across the NGF-induced differentiation time course (Day 0, Day 2, Day 4, and Day 6) were embedded using UMAP, revealing seven distinct clusters with separable protein expression patterns (left). Each cluster represents a group of cells sharing similar proteomic trajectories over time. (**B-F**) Line plots (right) show the relative protein abundance trends across differentiation days for proteins within each cluster, highlighting distinct temporal expression patterns across early, intermediate, and late stages.

Protein-level changes within selected clusters are shown in Figure 9. Cluster 1 contains proteins that are relatively abundant at the early stages of the time course and gradually decrease during differentiation, reaching their lowest levels around Day 4 before showing a modest recovery by Day 6. Enrichment analysis indicates that this cluster is dominated by proteins involved in ribosome structure, translation initiation, and RNA metabolism. Many of the most significant proteins belong to the ribosomal protein families (RPS and RPL), which form the core of the cellular translation machinery. The decline of these proteins over the time course suggests a reduction in global biosynthetic activity as cells transition away from a proliferative state. Early in differentiation, higher levels of protein synthesis support cellular growth and maintenance, whereas later stages involve a shift away from general protein production as the cells begin to adopt a neuronal phenotype.

**Figure 9.**
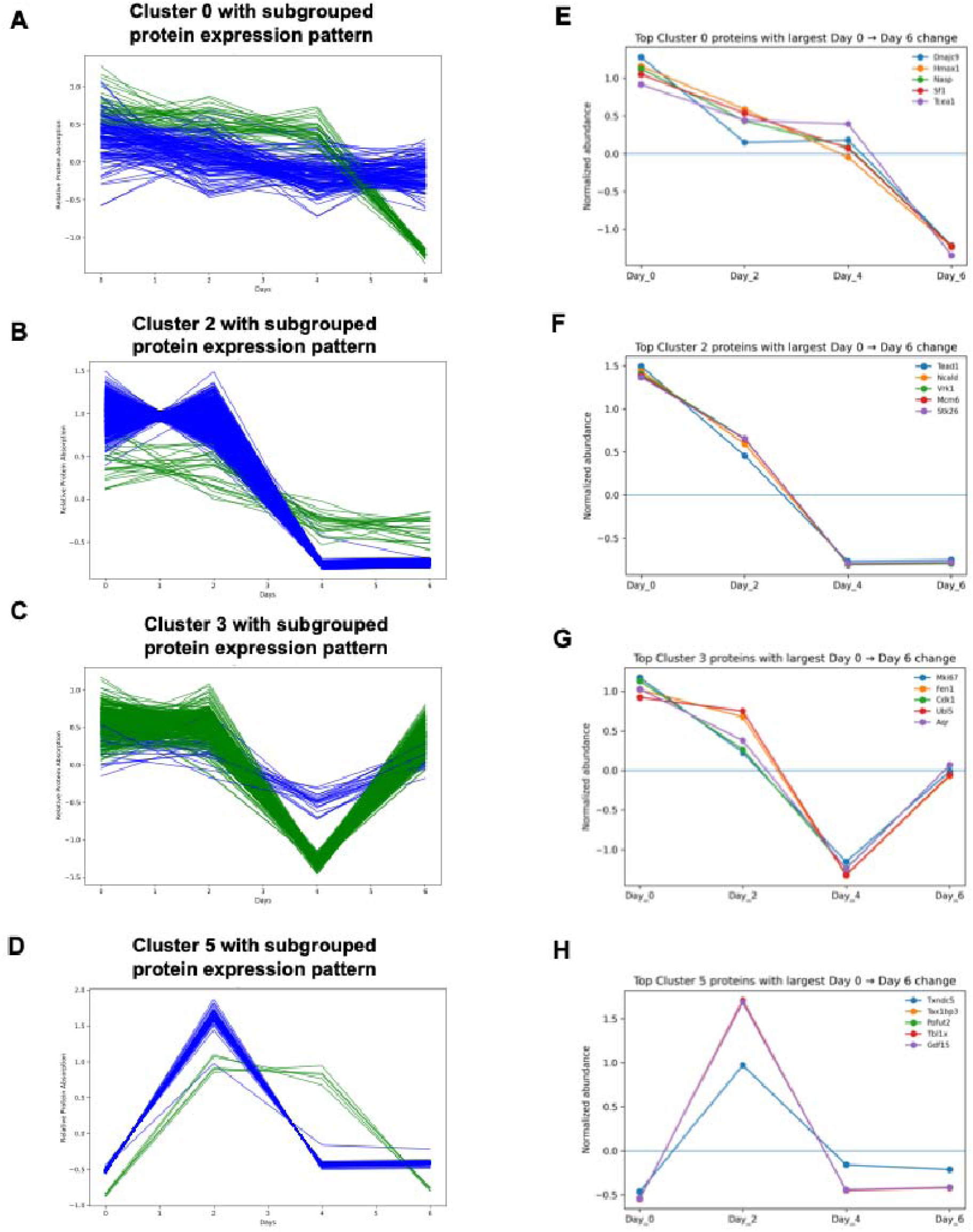
Subgrouped protein expression trajectories reveal additional heterogeneity within UMAP-defined clusters during PC12 differentiation. (A-D) Representative clusters (Clusters 0, 2, 3, and 5) were further resolved by separating proteins into distinct subgroups based on shared temporal expression patterns across the NGF-induced differentiation time course. **(E-H)** Line plots show relative protein abundance trajectories from Day 0 to Day 6, with proteins partitioned into two dominant expression subgroups (blue and green) within each cluster.

In contrast, Cluster 6 displays the opposite temporal trend, with proteins starting at lower abundance and progressively increasing toward the later stages of differentiation, particularly by Day 6. Pathway enrichment analysis shows strong representation of processes related to nucleocytoplasmic transport, RNA export, and mRNA processing. Several components of the nuclear transport and RNA trafficking machinery—including NXF1, NUP62, RAE1, RAN, and multiple RNA helicases—appear repeatedly among the most significant pathways. The increased abundance of these proteins during the later stages of differentiation likely reflects enhanced regulation of RNA trafficking and post-transcriptional control as neuronal programs become established. Efficient RNA transport between the nucleus and cytoplasm is critical for regulating gene expression and supporting localized protein synthesis, processes that are particularly important during neuronal maturation and structural remodeling. Together, these two clusters illustrate a coordinated molecular transition during NGF-induced differentiation, where early biosynthetic and translational programs give way to RNA regulatory processes associated with neuronal development.

For Clusters 0, 2, 3, and 5, proteins were ranked based on the absolute difference in normalized abundance between Day 0 and Day 6. Proteins with the largest changes were selected for display. In Cluster 0, several proteins show higher abundance at Day 0 followed by lower abundance at later time points, including Dnajc9, Hmox1, Nasp, Srf1, and Tcea1. In Cluster 2, proteins such as Tead1, Ncald, Vrk1, Mcm6, and Stk26 show reduced abundance by Day 4, with similar levels at Day 6.

In Cluster 3, proteins associated with cell cycle processes, including Mki67, Fen1, Cdk1, Ubl5, and Aqr, decrease between Day 2 and Day 4, followed by higher abundance at Day 6 relative to Day 4. In Cluster 5, proteins including Txndc5, Tax1bp3, Pofut2, Tbl1x, and Gdf15 show higher abundance at Day 2 compared with Day 0, with lower abundance at later time points. These cluster-level summaries show that proteins within UMAP-defined groups change across differentiation days, with different patterns observed across clusters.

Several proteins defining the temporal clusters are associated with stress response, transcription, and cell cycle regulation. Stress- and protein homeostasis–related proteins, including Dnajc9, Hmox1, and Txndc5, show higher abundance at early time points and decrease over the differentiation time course. Proteins involved in transcriptional regulation, such as Tcea1, Tead1, and Tbl1x, are more abundant at early or intermediate stages. In contrast, cell cycle–related proteins, including Mki67, Cdk1, Mcm6, Fen1, and Vrk1, decrease progressively with differentiation. These temporal trends are not evident from pairwise comparisons between individual time points (Ryter, S. W. *et al*. 2006; Wind, M., 2000; Zhao, B. *et al*., 2008; Scholzen, T., 2000; Malumbres, M, 2009).

### 4.13 Applied non-negative matrix factorization for cluster analysis and identification of proteins contributing to group separation

Application of NMF resolved three protein profiles that captured major sources of variation across the single-cell proteomics data (Figure S7A). When cells were projected into the space defined by these profiles, undifferentiated cells clustered within a narrow region characterized by higher contribution from profile 1. Cells collected at later differentiation stages occupied a broader region of profile space defined mainly by profiles 2 and 3.

At Days 4 and 6, cells separated into two partially overlapping groups distinguished by different combinations of profiles 2 and 3. By Day 6, most cells showed reduced contribution from profile 1. A subset of Day 6 cells also showed reduced contribution from profile 3, indicating additional variation within the late differentiation stage.

Inspection of the protein-by-profile loading matrix identified a limited number of proteins that contributed strongly to each profile (Figure S7B). These proteins differed across profiles and accounted for much of the observed separation between cell groups. Cells within the same differentiation stage often showed different profile combinations, indicating variability in proteomic state among cells sampled at the same time point.

#### 4.4.14 Proteins contributing to early subgroup separation at Day 2

Proteins contributing to the separation of the two Day 2 groups were examined by inspecting loadings along PC1 (Figure S8; Table S10). A subset of proteins showed relatively large positive or negative contributions to PC1, with z-scores close to ±2 and p-values below 0.05. These proteins accounted for much of the variance distinguishing the two groups at this time point.

Several of the highest-loading proteins are associated with translational and RNA-related processes, including Nat10, Eprs1, and Eif3f, along with proteins involved in intracellular trafficking such as Arl1. Proteins were distributed on both sides of PC1, indicating that the two Day 2 groups differ in opposing patterns of protein abundance rather than a shared directional change.

Proteins commonly used as markers of neuronal differentiation were not prominent among the PC1-associated features. Instead, the proteins contributing to group separation are consistent with differences in regulatory and biosynthetic activity, suggesting that variability at Day 2 reflects differences in cellular state rather than commitment to a neuronal differentiation program (Zismanov, V *et al*., 2016; Raj, B. *et al*., 2015).

#### 4.4.15 Proteins contributing to heterogeneity at Day 4 and Day 6

To further examine subgroup-specific proteomic features associated with divergent differentiation outcomes, we compared signaling and proteome remodeling patterns between Group A and Group B cells during NGF treatment (Figure 10). Group A cells showed higher abundance of endosomal trafficking proteins SNX1 and VPS26B, along with elevated levels of translational and proteostasis-related proteins (EEF1A2, HSP90AB1, SARNP, RPSA, and RPS21). These features were accompanied by increased abundance of cytoskeletal and neuronal maturation–associated proteins (TUBA1A, MAP1B, NEFL, NEFM, PRPH, and VGF) and coincided with neurite outgrowth and neuron-like phenotypes (Glock, C *et al*., 2017). In contrast, Group B cells exhibited lower abundance of proteins involved in receptor trafficking, translation, and cytoskeletal organization, aligning with weaker NGF-associated proteomic responses and the absence of neurite extension. VPS26B and SNX1 were selected as representative markers because their abundance reflects differences in endosomal trafficking capacity (Roundtree, I.A. *et al*., 2017).

**Figure 10.**
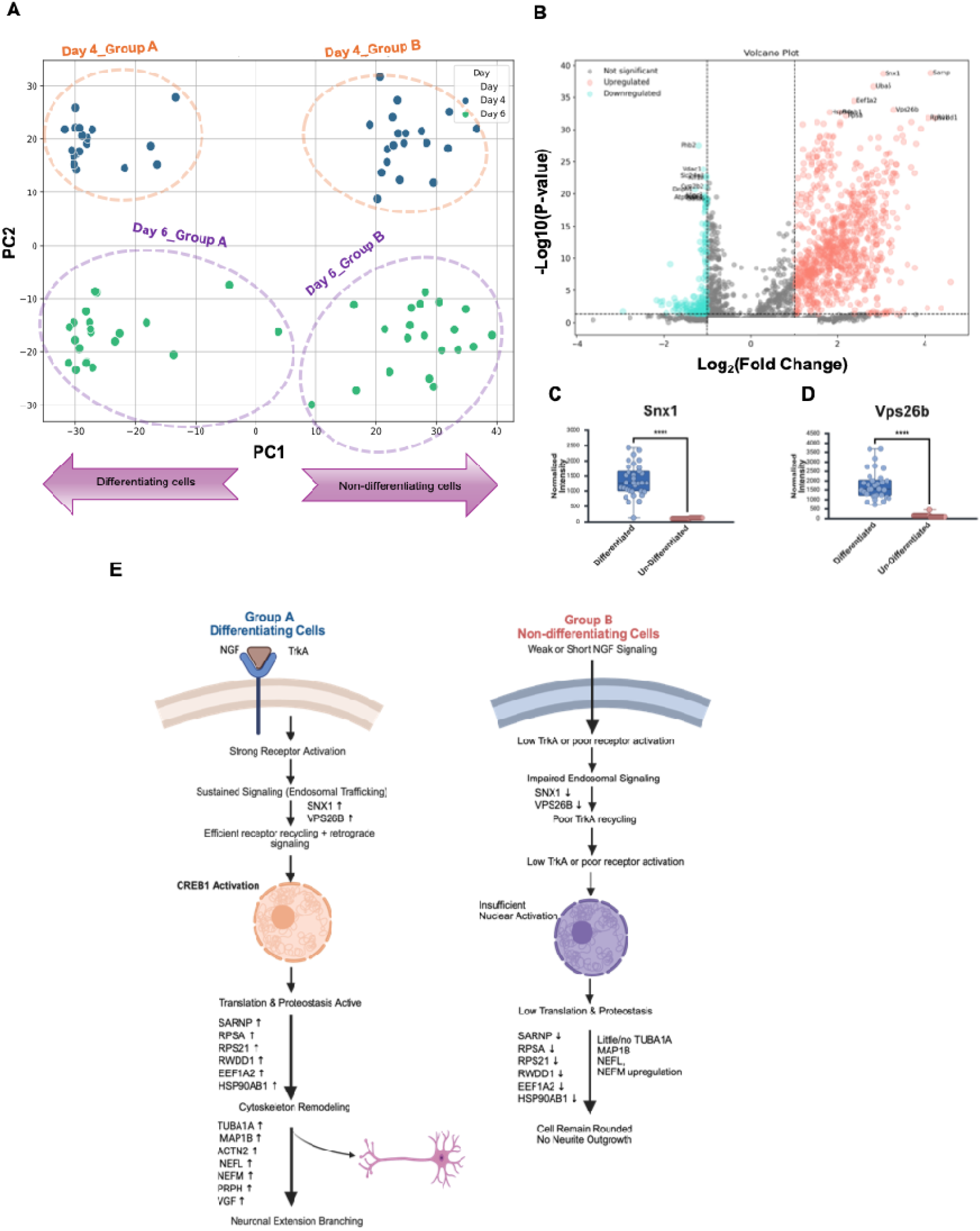
Subpopulation-specific proteomic remodeling during NGF-induced differentiation. **(A)** PCA plot showing separation of Group A and Group B subpopulations at Days 4 and 6. **(B)** Volcano plot of proteins differentially expressed between the two groups. |log FC| ≥ 1, p < 0.05). Proteins meeting both thresholds are highlighted. **(C-D)** Relative abundance of VPS26B and SNX1 in Group A and Group B cells. **(E)** Schematic summary of NGF-responsive pathways associated with the two subpopulations. Created with Biorender.com.

between subgroups: VPS26B was enriched in Group A cells, whereas SNX1 showed reduced abundance in Group B cells. Collectively, these subgroup-resolved proteomic profiles show how single-cell proteomics enables the identification of divergent molecular states within a heterogeneous population, differences that would be averaged and therefore masked in bulk proteomic measurements (Figure S9).

### 4.5 Discussion

Both morphological assessment and proteomic profiling show that PC12 differentiation does not occur uniformly across the cell population. While many cells exhibit neurite outgrowth and neuron-like features by Days 4 and 6, others remain partially or fully undifferentiated. This variability is reflected in the proteomic data, where later time points show increased dispersion in PCA, multimodal protein abundance distributions, and separation into multiple subclusters. In contrast, Day 0 cells cluster more tightly and align with a relatively homogeneous proliferative state. NGF-induced differentiation did not progress uniformly across the cell population, with individual cells occupying multiple proteomic states within the same differentiation stage.

Capturing this heterogeneity required careful optimization of cell handling and sample preparation. Differentiated PC12 cells are larger, more fragile, and prone to aggregation, making gentle dissociation, anti-clumping strategies, and careful tuning of the dispensing parameters essential. These optimizations enabled high single-cell dispensing accuracy and reproducibility, providing consistent input for DIA-based proteomic analysis. Quality control metrics confirmed stable quantitative performance across all time points, with similar precursor-level variability and overlapping intensity distributions. Although fewer proteins were identified at later stages, this likely reflects biological changes associated with differentiation, including cell-cycle exit, reduced global protein synthesis, and increased membrane specialization, rather than technical limitations alone. The improved protein recovery observed with DDM further emphasizes the importance of optimized extraction strategies for low-input and differentiated cell types.

Beyond revealing cell-to-cell heterogeneity, single-cell proteomics provides insight into regulatory mechanisms that are inherently obscured in bulk measurements, particularly during dynamic biological processes such as neuronal differentiation. Differentiation involves asynchronous transitions, transient signaling events, and stage-specific remodeling of cytoskeletal, trafficking, and metabolic pathways. Bulk proteomics captures the mean signal across thousands of cells, which can mask transitional states and compress dynamic range, especially for proteins that are tightly regulated or expressed in a subset of cells. In contrast, single-cell approaches preserve the distribution of protein abundance across individual cells, enabling detection of rare states, asynchronous differentiation trajectories, and coordinated remodeling within specific subpopulations (Specht and Slavov, 2018; Budnik *et al*., 2018). This is particularly relevant in neuronal systems, where neurite extension, synaptic maturation, and axonal transport are not uniformly activated across all cells at the same time. Moreover, protein-level regulation often diverges from transcript-level patterns due to post-transcriptional control, differences in translation efficiency, and protein stability (Liu, Beyer, and Aebersold, 2016). Therefore, single-cell proteomics not only refines our understanding of heterogeneity but also captures regulatory complexity at the functional protein level, providing a more accurate representation of the molecular programs governing neuronal growth and maturation.

In summary, this work demonstrates the utility of single-cell proteomics for studying dynamic and heterogeneous biological processes such as neuronal differentiation. This workflow enabled reproducible protein measurements across multiple days of differentiation and captured intermediate proteomic features that are not resolved in bulk analyses. Although current single-cell proteomics platforms remain limited in throughput, incremental improvements in sample preparation, chromatographic separation, and mass spectrometry are expected to enhance performance. Applying similar approaches to primary neuronal cultures or cells derived from in vivo models and pairing proteomic measurements with complementary assays such as imaging or transcript profiling, may further clarify molecular changes associated with neuronal differentiation and maturation.

## Author Contributions

Arpa Ebrahimi performed the experiments, collected and processed the data, interpreted the results, and wrote the manuscript. Shuxin Chi conducted experiments, contributed to data collection and processing, and assisted with manuscript review and editing. Jun Yang performed bioinformatics analyses and contributed to data interpretation, manuscript review, and editing. Phoebe Y. Lee assisted with experimental procedures and data analysis and contributed to manuscript review and editing. Prongbaramee K. Colling assisted with experimental procedures and contributed to manuscript review and editing. Leonard J. Foster provided guidance on study design, contributed to data interpretation, and reviewed and edited the manuscript. David A. Hendrix provided computational advice and contributed to manuscript review and editing. Claudia S. Maier supervised the project, guided experimental design and data interpretation, and contributed to manuscript review and editing. All authors approved the final version of the manuscript.

## Acknowledgments

We thank the University of British Columbia Proteomics Core Facility for access to the instrument, technical support, and for their collaboration throughout this project. We also acknowledge Dr. Fredrick Stevens for providing access to the cell culture facility and for technical assistance. We are grateful to HP Lifesciences, especially Dr. Hyo Sang Jang, for their collaboration and support in developing and optimizing single-cell proteomics workflows. Finally, we thank the members of the Maier laboratory for their helpful discussions and feedback throughout this work.

## Data Availability Statement

The raw single-cell proteomics mass spectrometry data generated in this study have been deposited in the MassIVE repository under accession number **MSV000100666**.

---

**Figure S1.**
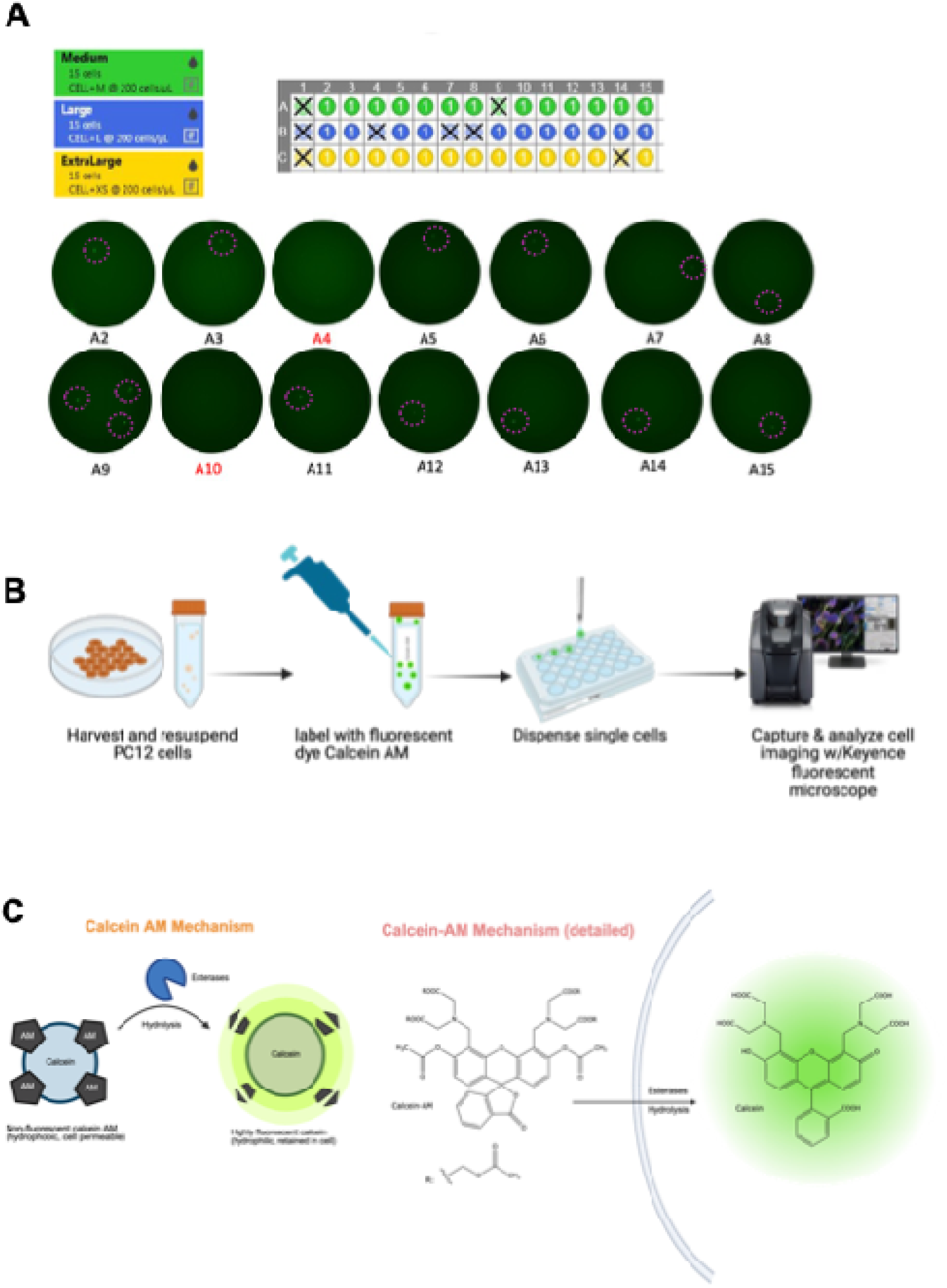
Optimization and validation of single-cell dispensing for neuronal cells using the HP D100 platform. **(A)** Single cell dispenser software interface and optimization of the Medium, Large, and X-Large dispensing settings to accommodate the size and morphology of neuronal cells. **(B)** Calcein AM fluorescence microscopy was used to verify the accuracy of single-cell dispensing and assess the viability of post-dispensed cells. Fluorescent labeling confirms both proper deposition of individual cells into wells and retention of membrane integrity following dispensing. **(C)** Schematic of the Calcein AM labeling mechanism, in which the non-fluorescent, cell-permeable dye is converted by intracellular esterases into fluorescent Calcein, enabling visualization of viable cells.

**Figure S2.**
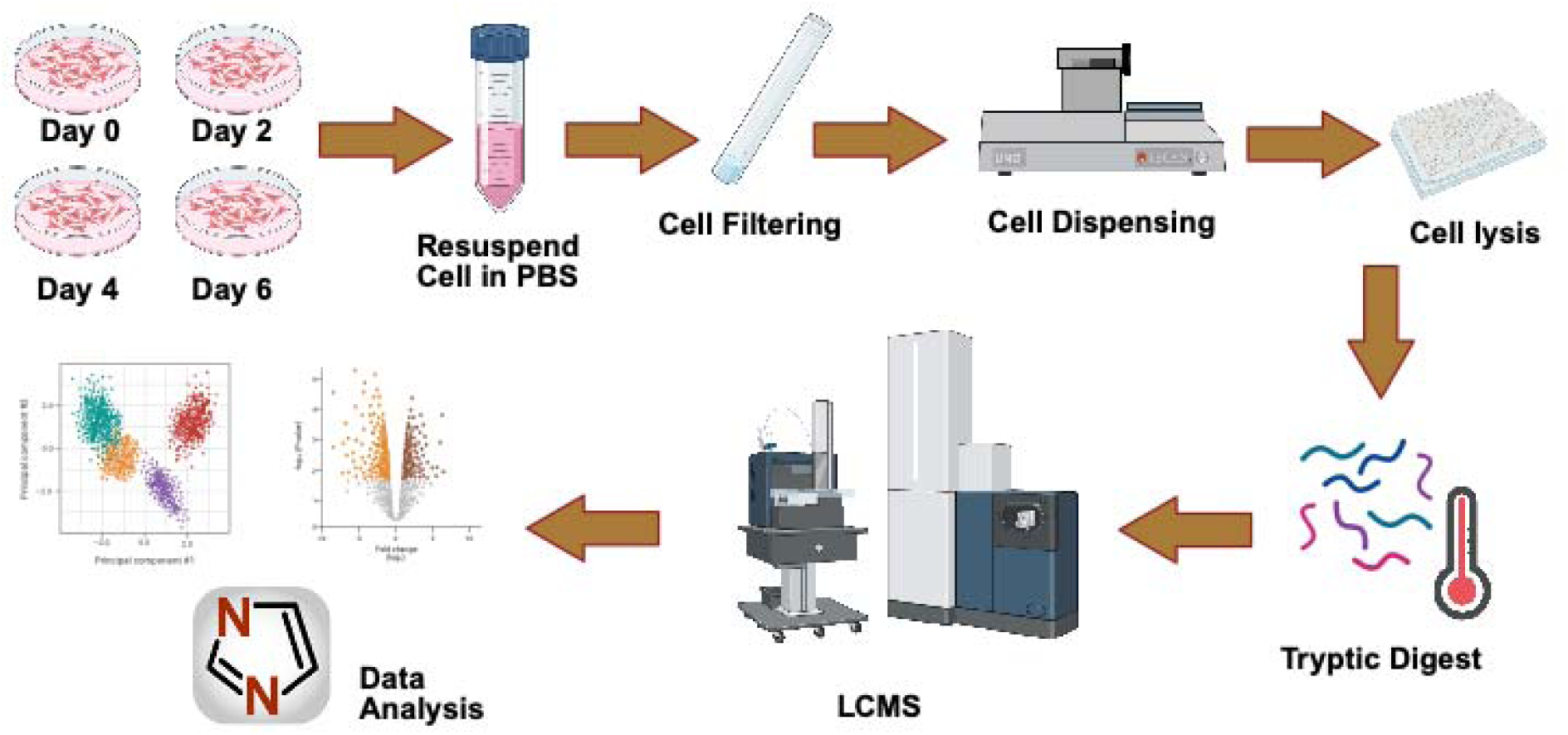
Single-cell proteomic sample preparation and proteomics workflow for PC12 cells. PC12 cells were cultured, treated with NGF, and harvested at defined time points. Single cells were isolated, lysed, and digested with trypsin before LC-MS/MS analysis and dia-PASEF data processing. Data were processed for peptide identification, quantification, and statistical comparison across differentiation stages. Created with Biorender.com.

**Figure S3.**
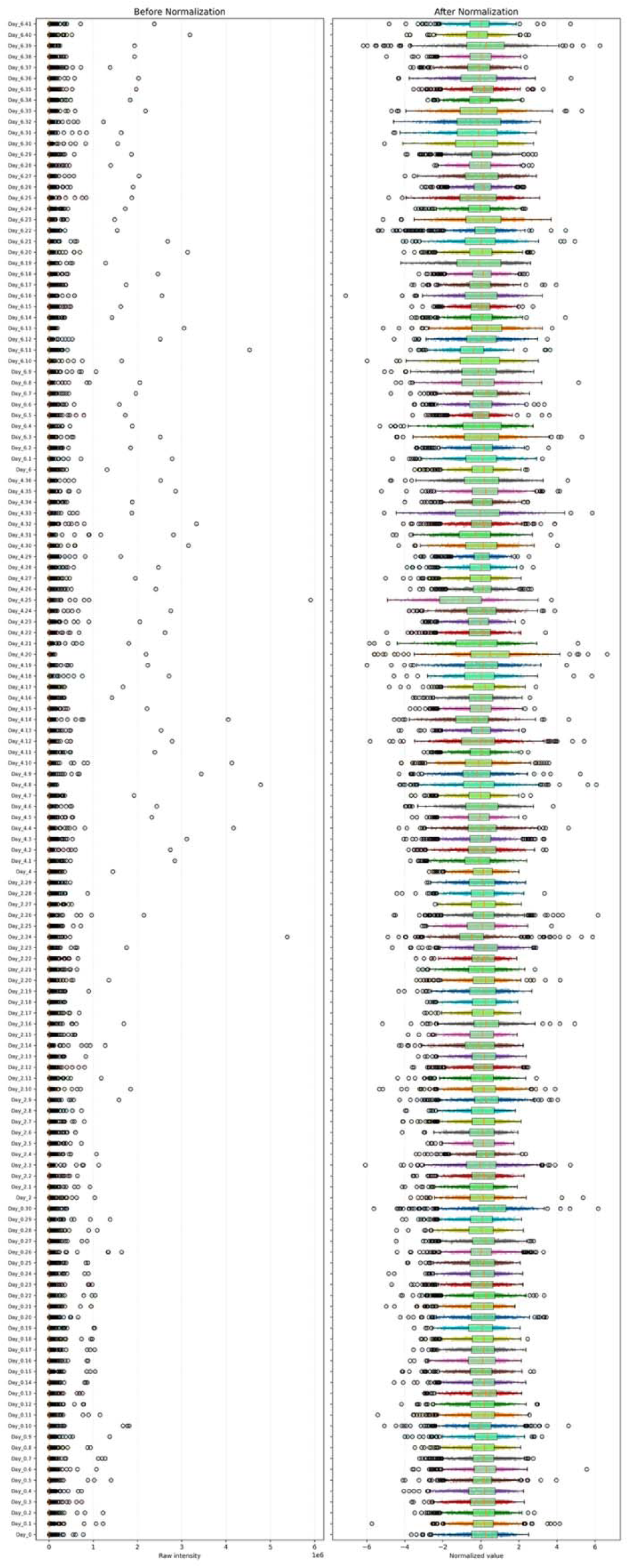
Total-sum normalization of protein intensities. Protein intensity distributions across samples are shown before and after normalization. To account for differences in total signal between samples, protein intensities were scaled so that each sample had the same total intensity. Following normalization, the distributions are comparable across samples, indicating that global intensity differences were effectively corrected prior to downstream quantitative analysis.

**Figure S4.**
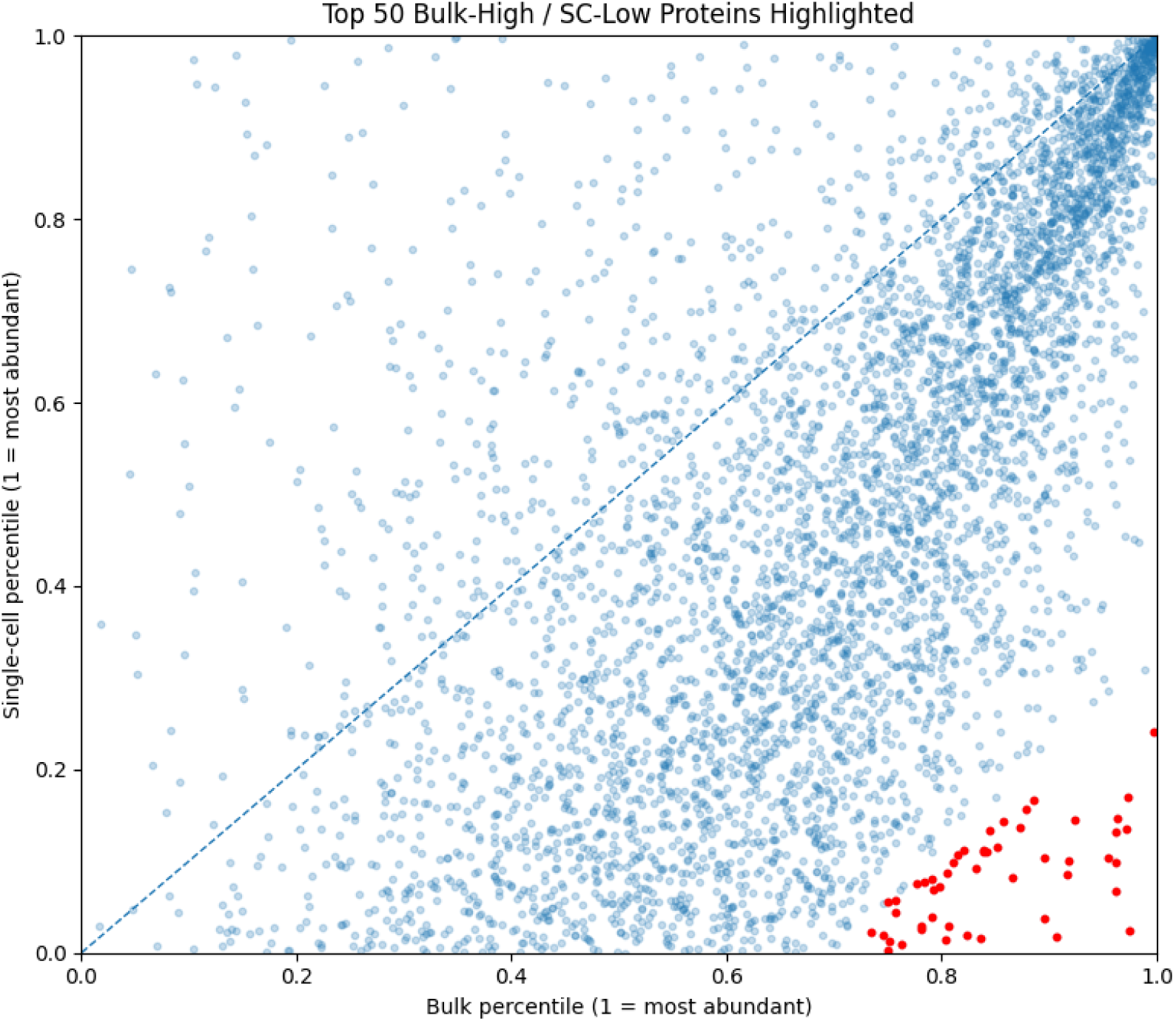
Percentile-based comparison of bulk and single-cell protein abundance rankings. Protein abundance ranks from bulk and single-cell datasets were independently converted to percentile scores using percentile = 1 − ((rank − 1)/N), where N is the total number of detected proteins in each dataset. This normalization rescales ranks from 0 to 1 (1 = highest relative abundance) and enables direct comparison despite differences in proteome depth. Percentiles for proteins detected in both datasets were plotted with bulk values on the x-axis and single-cell values on the y-axis. The dashed line (y = x) indicates equal relative abundance between modalities. Proteins deviating from the identity line reflect discordant abundance patterns, and the top 50 proteins with the largest positive Δ(Bulk − Single-cell percentile) are highlighted to illustrate proteins that appear highly abundant in bulk but relatively low in single-cell measurement

**Figure S5.**
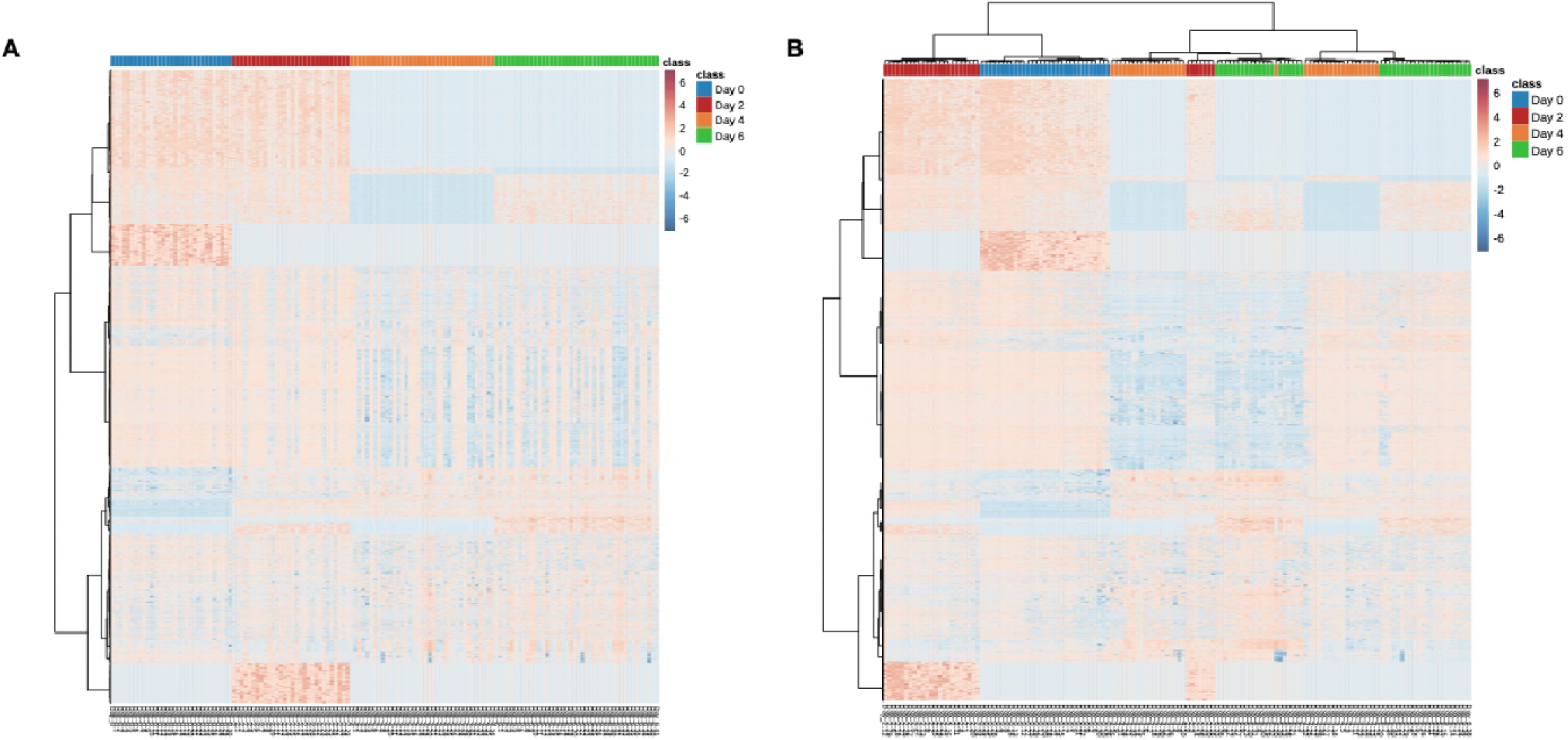
Heatmaps of single-cell protein abundance across differentiation stages shown without (A) and with (B) sample clustering. **(A)** In the unclustered heatmap, samples are ordered by differentiation time point (Day 0, Day 2, Day 4, Day 6), emphasizing average temporal trends. **(B)** In the clustered heatmap, samples are reordered based on similarity in protein abundance patterns, revealing distinct subpopulations within Day 2, Day 4, and Day 6 characterized by opposing protein expression profiles. Color scale represents z-scored protein abundance.

**Figure S6.**
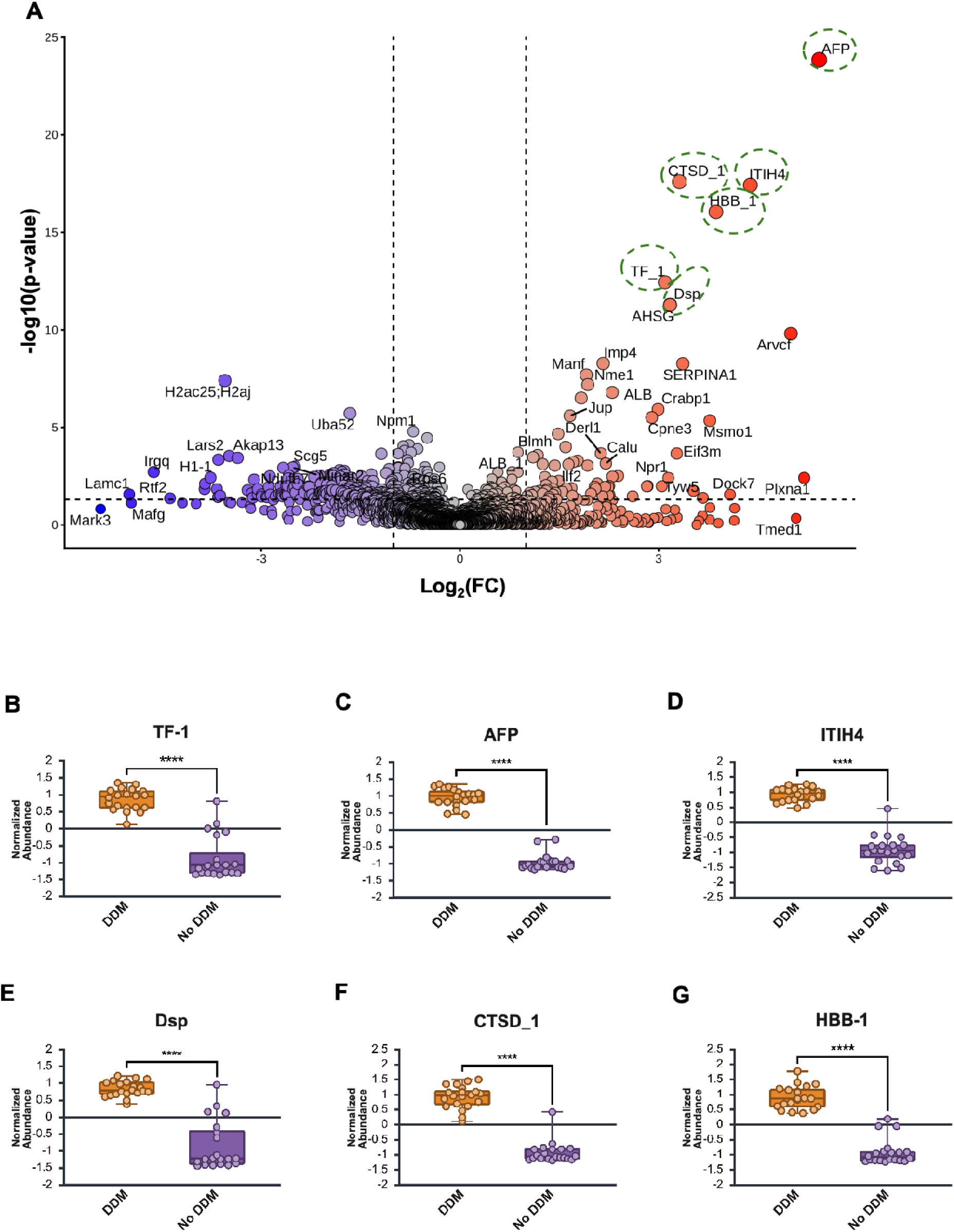
Effect of DDM on insoluble and membrane protein recovery in differentiated neuronal cells. **(A)** Volcano plot comparing Day 6 differentiated neuronal cells prepared with and without n-dodecyl-β-D-maltoside (DDM). Proteins with a fold change >2 and *p* < 0.05 are highlighted. **(B–G)** Bar graphs showing representative proteins significantly enriched in DDM-treated samples, illustrating enhanced recovery of vesicle-associated, membrane-proximal, and poorly soluble proteins. Statistical significance was assessed using a Mann–Whitney U test; **** denotes *p* < 0.0001.

**Figure S7.**
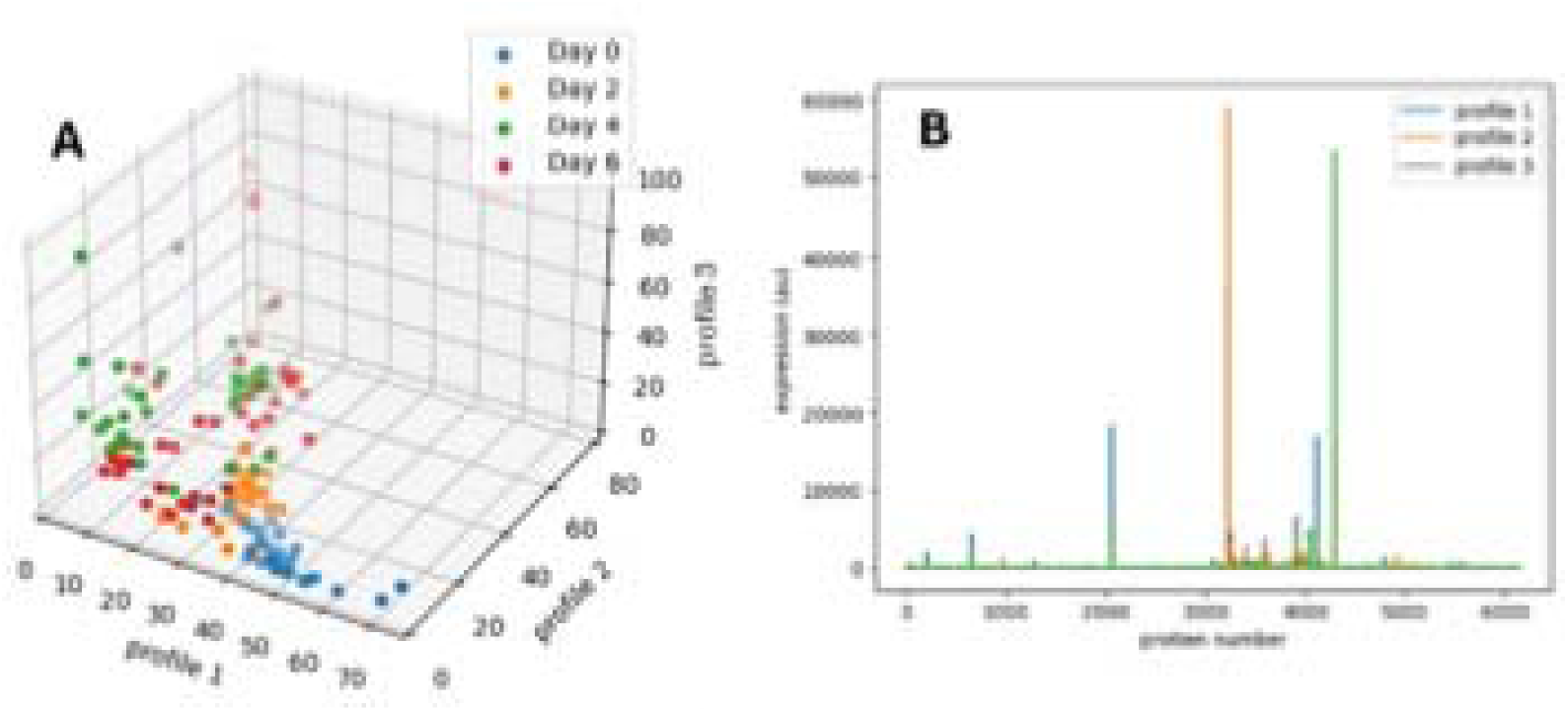
Applied non-negative matrix factorization for cluster analysis and identification of proteins contributing to group separation. **(A)** Points represent individual cells projected into a three-profile space derived from non-negative matrix factorization. Axes correspond to the relative expression of protein profiles that combine to approximate total protein abundance in each cell. Cells are colored by differentiation time point. **(B)** Protein loadings for each profile, showing proteins that contribute to the separation between cell groups.

**Figure S8.**
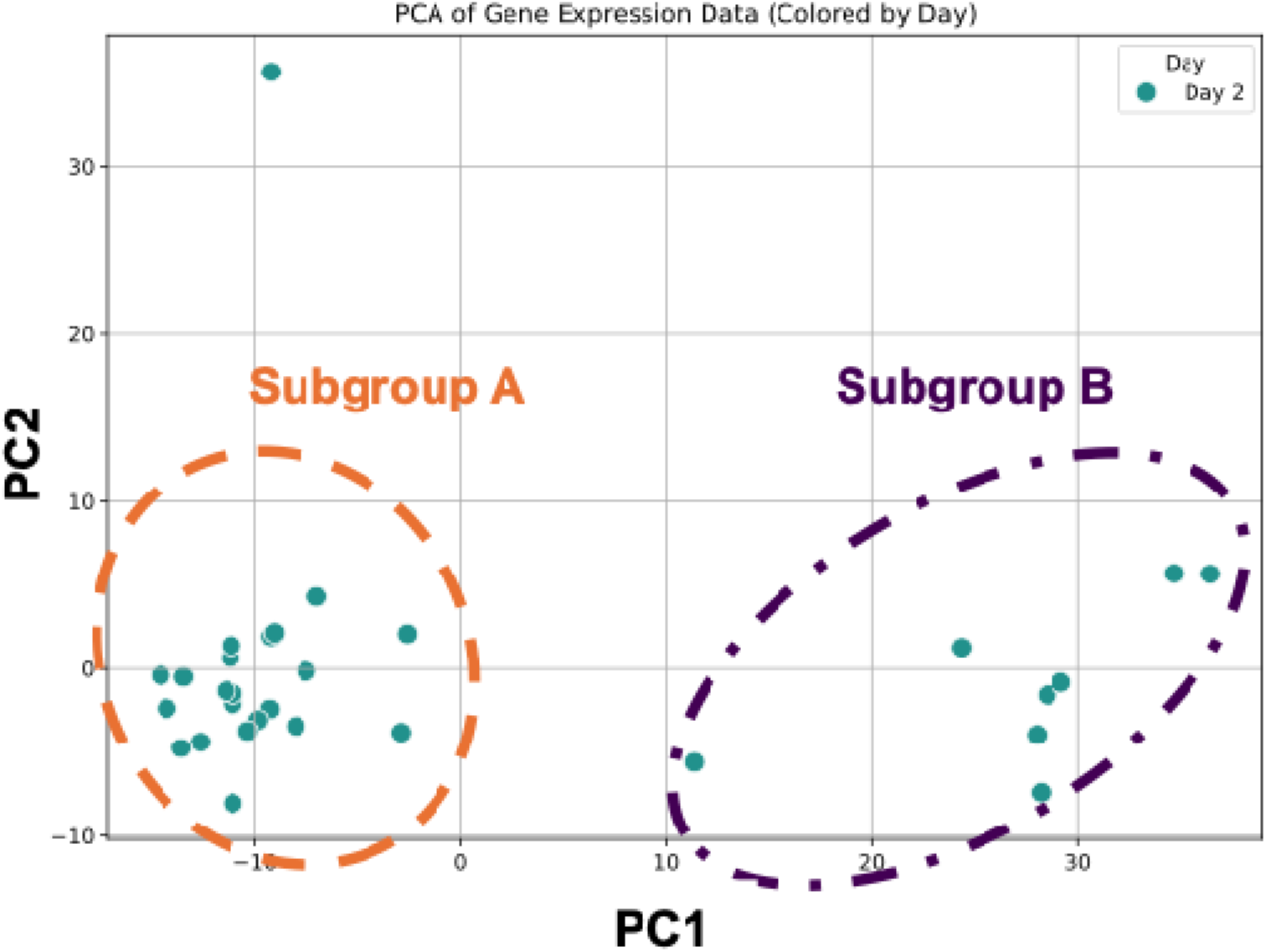
Principal component analysis of single-cell proteomic profiles at Day 2 resolves two distinct subgroups primarily along PC1.

**Figure S9.**
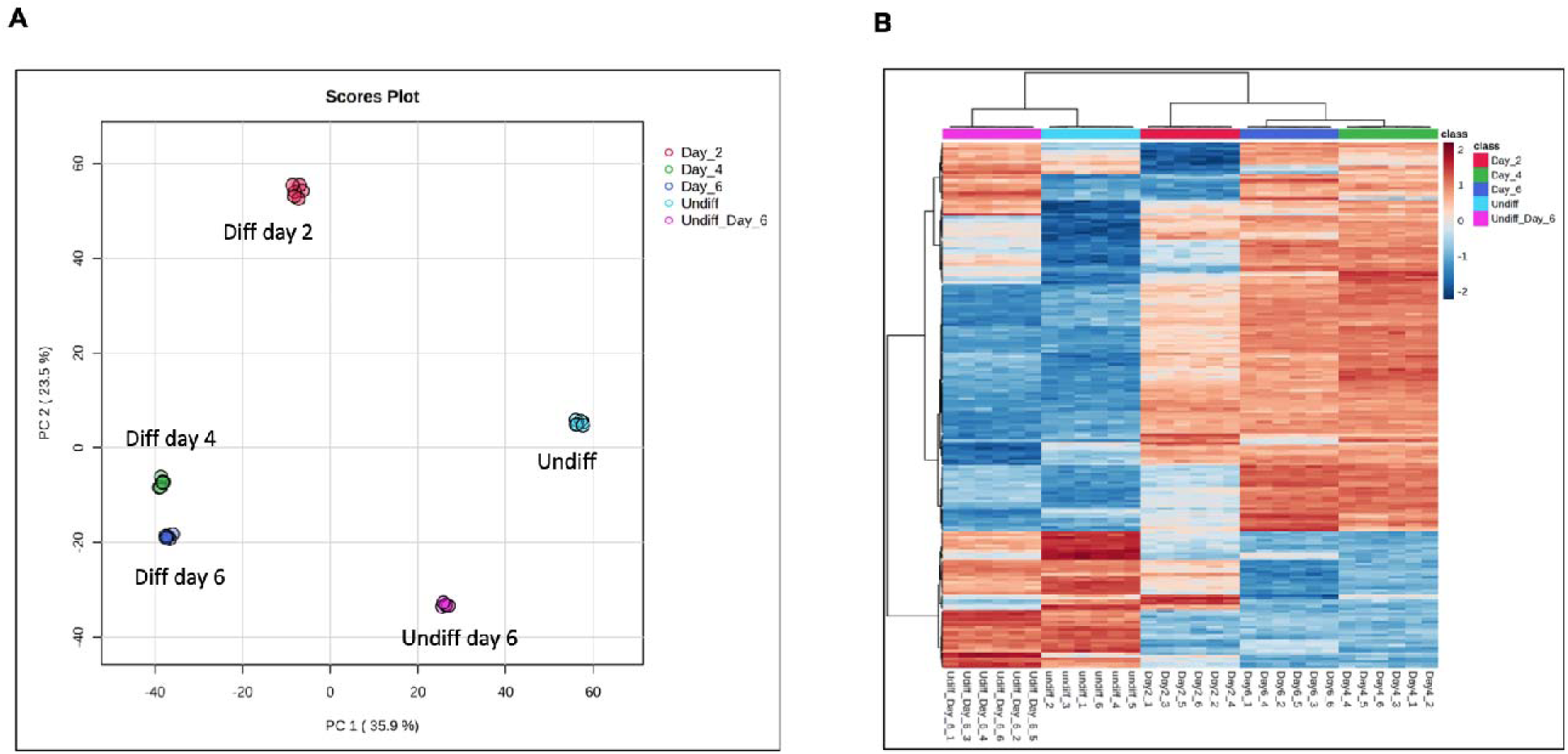
Bulk proteomics analysis of PC12 neuronal differentiation. **(A)** Principal component analysis (PCA) of bulk proteomic profiles collected at Day 0, Day 2, Day 4, and Day 6 following NGF treatment (n = 5 analytical replicates per time point). PCA shows clear separation between early Day 0, Day 2, and later differentiation stages (Day 4–Day 6), indicating reproducible, time-dependent proteome remodeling at the population level. Replicates within each time point cluster tightly, reflecting low technical variability in the bulk measurements. **(B)** Heatmap of bulk protein abundance across differentiation stages. Protein intensities were log-transformed and z-score normalized across samples. The heatmap highlights systematic changes in protein abundance between time points, with distinct expression patterns emerging at Day 2, Day 4, and Day 6. While bulk analysis captures robust temporal trends in proteome remodeling, the tight clustering of replicates and absence of within-group structure indicate that bulk measurements average across cells and do not resolve the cell-to-cell heterogeneity observed in the single-cell proteomics data.

**Table S1.** HP D100 dispenser software settings for reagent dispensing.

| Parameter | Software setting |
| --- | --- |
| Dispensing mode | Dispense by volume |
| Fluid group | Aqueous |
| Fluid class | Aqueous (surfactant-free) |
| Dispense volume | 1 μL |
| Set value | 1 μL per well |
| Dispense pattern | Single dispense per well |
| Plate format | 384-well plate |
| Control software | HP D100 Control Software (v3.7.0) |

**Table S2.** HP D100 dispenser software settings for single-cell dispensing.

| Parameter | Software setting |
| --- | --- |
| Dispensing mode | Cell dispensing |
| Fluid group | Sticky Cells |
| Fluid class | Medium <sup>1</sup> |
| Cell concentration | 200 cells/μL |
| Set value | 1, 5, or 10 cells per well<sup>2</sup> |
| Dispense pattern | Single dispense per well |
| Plate format | 384-well plate |
| Control software | HP D100 Control Software (v3.7.0) |

**Table S3.** Single-cell proteomics sample metadata. This table lists metadata for all single-cell PC12 samples analyzed, including sample identifiers, differentiation time point, acquisition date, 384-well plate position, and raw data file names. Samples include undifferentiated (Day 0; N = 35) and NGF-differentiated PC12 cells collected at Day 2 (N = 36), Day 4 (N = 42), and Day 6 (N = 50). Blank injections and 250 pg HeLa digest quality control samples were acquired after every 10 single-cell sample injections to monitor system performance and carryover.

| Sample ID | Cell type | Time point | Acquisition date | 384-well plate position | Sequence name |
| --- | --- | --- | --- | --- | --- |
| SC_PC12_D0_01 | PC12 | Day 0 | 8/26/24 | 27 | Arpa_PC12_day0_8-26-2024_Slot1-27_7583.d |
| SC_PC12_D0_02 | PC12 | Day 0 | 8/26/24 | 28 | Arpa_PC12_day0_8-26-2024_Slot1-28_7583.d |
| SC_PC12_D0_03 | PC12 | Day 0 | 8/26/24 | 29 | Arpa_PC12_day0_8-26-2024_Slot1-29_7584.d |
| SC_PC12_D0_04 | PC12 | Day 0 | 8/26/24 | 30 | Arpa_PC12_day0_8-26-2024_Slot1-30_7585.d |
| SC_PC12_D0_05 | PC12 | Day 0 | 8/26/24 | 31 | Arpa_PC12_day0_8-26-2024_Slot1-31_7586.d |
| SC_PC12_D0_06 | PC12 | Day 0 | 8/26/24 | 32 | Arpa_PC12_day0_8-26-2024_Slot1-32_7587.d |
| SC_PC12_D0_07 | PC12 | Day 0 | 8/27/24 | 33 | Arpa_PC12_day0_8-27-2024_Slot1-33_7588.d |
| SC_PC12_D0_08 | PC12 | Day 0 | 8/27/24 | 34 | Arpa_PC12_day0_8-27-2024_Slot1-34_7589.d |
| SC_PC12_D0_09 | PC12 | Day 0 | 8/27/24 | 35 | Arpa_PC12_day0_8-27-2024_Slot1-35_7590.d |
| SC_PC12_D0_10 | PC12 | Day 0 | 8/27/24 | 36 | Arpa_PC12_day0_8-27-2024_Slot1-36_7591.d |
| SC_PC12_D0_11 | PC12 | Day 0 | 8/27/24 | 37 | Arpa_PC12_day0_8-27-2024_Slot1-37_7593.d |
| SC_PC12_D0_12 | PC12 | Day 0 | 8/27/24 | 38 | Arpa_PC12_day0_8-27-2024_Slot1-38_7594.d |
| SC_PC12_D0_13 | PC12 | Day 0 | 8/27/24 | 39 | Arpa_PC12_day0_8-27-2024_Slot1-39_7595.d |
| SC_PC12_D0_14 | PC12 | Day 0 | 8/27/24 | 40 | Arpa_PC12_day0_8-27-2024_Slot1-40_7596.d |
| SC_PC12_D0_15 | PC12 | Day 0 | 8/27/24 | 41 | Arpa_PC12_day0_8-27-2024_Slot1-41_7597.d |
| SC_PC12_D0_16 | PC12 | Day 0 | 8/27/24 | 42 | Arpa_PC12_day0_8-27-2024_Slot1-42_7598.d |
| SC_PC12_D0_17 | PC12 | Day 0 | 8/27/24 | 43 | Arpa_PC12_day0_8-27-2024_Slot1-43_7599.d |
| SC_PC12_D0_18 | PC12 | Day 0 | 8/27/24 | 44 | Arpa_PC12_day0_8-27-2024_Slot1-44_7600.d |
| SC_PC12_D0_19 | PC12 | Day 0 | 8/27/24 | 45 | Arpa_PC12_day0_8-27-2024_Slot1-45_7601.d |
| SC_PC12_D0_20 | PC12 | Day 0 | 8/27/24 | 46 | Arpa_PC12_day0_8-27-2024_Slot1-46_7602.d |
| SC_PC12_D0_21 | PC12 | Day 0 | 8/27/24 | 52 | Arpa_PC12_day0_8-27-2024_Slot1-52_7604.d |
| SC_PC12_D0_22 | PC12 | Day 0 | 8/27/24 | 53 | Arpa_PC12_day0_8-27-2024_Slot1-53_7605.d |
| SC_PC12_D0_23 | PC12 | Day 0 | 8/27/24 | 54 | Arpa_PC12_day0_8-27-2024_Slot1-54_7606.d |
| SC_PC12_D0_24 | PC12 | Day 0 | 8/27/24 | 55 | Arpa_PC12_day0_8-27-2024_Slot1-55_7607.d |
| SC_PC12_D0_25 | PC12 | Day 0 | 8/27/24 | 56 | Arpa_PC12_day0_8-27-2024_Slot1-56_7608.d |
| SC_PC12_D0_26 | PC12 | Day 0 | 8/27/24 | 57 | Arpa_PC12_day0_8-27-2024_Slot1-57_7609.d |
| SC_PC12_D0_27 | PC12 | Day 0 | 8/27/24 | 58 | Arpa_PC12_day0_8-27-2024_Slot1-58_7610.d |
| SC_PC12_D0_28 | PC12 | Day 0 | 8/27/24 | 59 | Arpa_PC12_day0_8-27-2024_Slot1-59_7611.d |
| SC_PC12_D0_29 | PC12 | Day 0 | 8/28/24 | 60 | Arpa_PC12_day0_8-28-2024_Slot1-60_7612.d |
| SC_PC12_D0_30 | PC12 | Day 0 | 8/28/24 | 61 | Arpa_PC12_day0_8-28-2024_Slot1-61_7613.d |
| SC_PC12_D0_31 | PC12 | Day 0 | 8/28/24 | 85 | Arpa_PC12_day0_8-28-2024_Slot1-85_7614.d |
| SC_PC12_D0_32 | PC12 | Day 0 | 8/28/24 | 86 | Arpa_PC12_day0_8-28-2024_Slot1-86_7615.d |
| SC_PC12_D0_33 | PC12 | Day 0 | 8/28/24 | 87 | Arpa_PC12_day0_8-28-2024_Slot1-87_7616.d |
| SC_PC12_D0_34 | PC12 | Day 0 | 8/28/24 | 88 | Arpa_PC12_day0_8-28-2024_Slot1-88_7617.d |
| SC_PC12_D0_35 | PC12 | Day 0 | 8/28/24 | 89 | Arpa_PC12_day0_8-28-2024_Slot1-89_7618.d |
| SC_PC12_D2_01 | PC12 | Day 2 | 8/29/24 | 27 | Arpa_PC12_day2_8-29-2024_Slot1-27_7658.d |
| SC_PC12_D2_02 | PC12 | Day 2 | 8/29/24 | 28 | Arpa_PC12_day2_8-29-2024_Slot1-28_7659.d |
| SC_PC12_D2_03 | PC12 | Day 2 | 8/29/24 | 29 | Arpa_PC12_day2_8-29-2024_Slot1-29_7660.d |
| SC_PC12_D2_04 | PC12 | Day 2 | 8/29/24 | 30 | Arpa_PC12_day2_8-29-2024_Slot1-30_7661.d |
| SC_PC12_D2_05 | PC12 | Day 2 | 8/30/24 | 31 | Arpa_PC12_day2_8-30-2024_Slot1-31_7662.d |
| SC_PC12_D2_06 | PC12 | Day 2 | 8/30/24 | 32 | Arpa_PC12_day2_8-30-2024_Slot1-32_7663.d |
| SC_PC12_D2_07 | PC12 | Day 2 | 8/30/24 | 33 | Arpa_PC12_day2_8-30-2024_Slot1-33_7664.d |
| SC_PC12_D2_08 | PC12 | Day 2 | 8/30/24 | 34 | Arpa_PC12_day2_8-30-2024_Slot1-34_7665.d |
| SC_PC12_D2_09 | PC12 | Day 2 | 8/30/24 | 35 | Arpa_PC12_day2_8-30-2024_Slot1-35_7666.d |
| SC_PC12_D2_10 | PC12 | Day 2 | 8/30/24 | 36 | Arpa_PC12_day2_8-30-2024_Slot1-36_7667.d |
| SC_PC12_D2_11 | PC12 | Day 2 | 8/30/24 | 39 | Arpa_PC12_day2_8-30-2024_Slot1-39_7670.d |
| SC_PC12_D2_12 | PC12 | Day 2 | 8/30/24 | 40 | Arpa_PC12_day2_8-30-2024_Slot1-40_7671.d |
| SC_PC12_D2_13 | PC12 | Day 2 | 8/30/24 | 41 | Arpa_PC12_day2_8-30-2024_Slot1-41_7672.d |
| SC_PC12_D2_14 | PC12 | Day 2 | 8/30/24 | 42 | Arpa_PC12_day2_8-30-2024_Slot1-42_7676.d |
| SC_PC12_D2_15 | PC12 | Day 2 | 8/30/24 | 43 | Arpa_PC12_day2_8-30-2024_Slot1-43_7677.d |
| SC_PC12_D2_16 | PC12 | Day 2 | 8/30/24 | 44 | Arpa_PC12_day2_8-30-2024_Slot1-44_7678.d |
| SC_PC12_D2_17 | PC12 | Day 2 | 8/30/24 | 45 | Arpa_PC12_day2_8-30-2024_Slot1-45_7679.d |
| SC_PC12_D2_18 | PC12 | Day 2 | 8/30/24 | 46 | Arpa_PC12_day2_8-30-2024_Slot1-46_7680.d |
| SC_PC12_D2_19 | PC12 | Day 2 | 8/30/24 | 47 | Arpa_PC12_day2_8-30-2024_Slot1-47_7681.d |
| SC_PC12_D2_20 | PC12 | Day 2 | 8/30/24 | 50 | Arpa_PC12_day2_8-30-2024_Slot1-50_7682.d |
| SC_PC12_D2_21 | PC12 | Day 2 | 8/30/24 | 51 | Arpa_PC12_day2_8-30-2024_Slot1-51_7683.d |
| SC_PC12_D2_22 | PC12 | Day 2 | 8/30/24 | 52 | Arpa_PC12_day2_8-30-2024_Slot1-52_7684.d |
| SC_PC12_D2_23 | PC12 | Day 2 | 8/30/24 | 53 | Arpa_PC12_day2_8-30-2024_Slot1-53_7685.d |
| SC_PC12_D2_24 | PC12 | Day 2 | 8/30/24 | 54 | Arpa_PC12_day2_8-30-2024_Slot1-54_7686.d |
| SC_PC12_D2_25 | PC12 | Day 2 | 8/31/24 | 55 | Arpa_PC12_day2_8-31-2024_Slot1-55_7687.d |
| SC_PC12_D2_26 | PC12 | Day 2 | 8/31/24 | 56 | Arpa_PC12_day2_8-31-2024_Slot1-56_7688.d |
| SC_PC12_D2_27 | PC12 | Day 2 | 8/31/24 | 57 | Arpa_PC12_day2_8-31-2024_Slot1-57_7689.d |
| SC_PC12_D2_28 | PC12 | Day 2 | 8/31/24 | 58 | Arpa_PC12_day2_8-31-2024_Slot1-58_7690.d |
| SC_PC12_D2_29 | PC12 | Day 2 | 8/31/24 | 60 | Arpa_PC12_day2_8-31-2024_Slot1-60_7691.d |
| SC_PC12_D2_30 | PC12 | Day 2 | 8/31/24 | 61 | Arpa_PC12_day2_8-31-2024_Slot1-61_7693.d |
| SC_PC12_D2_31 | PC12 | Day 2 | 8/31/24 | 62 | Arpa_PC12_day2_8-31-2024_Slot1-62_7694.d |
| SC_PC12_D2_32 | PC12 | Day 2 | 8/31/24 | 63 | Arpa_PC12_day2_8-31-2024_Slot1-63_7695.d |
| SC_PC12_D2_33 | PC12 | Day 2 | 8/31/24 | 64 | Arpa_PC12_day2_8-31-2024_Slot1-64_7696.d |
| SC_PC12_D2_34 | PC12 | Day 2 | 8/31/24 | 65 | Arpa_PC12_day2_8-31-2024_Slot1-65_7697.d |
| SC_PC12_D2_35 | PC12 | Day 2 | 8/31/24 | 66 | Arpa_PC12_day2_8-31-2024_Slot1-66_7698.d |
| SC_PC12_D2_36 | PC12 | Day 2 | 8/31/24 | 67 | Arpa_PC12_day2_8-31-2024_Slot1-67_7699.d |
| SC_PC12_D4_1 | PC12 | Day 4 | 8/31/24 | 30 | Arpa_PC12_day4_8-31-2024_Slot1-30_7703.d |
| SC_PC12_D4_2 | PC12 | Day 4 | 8/31/24 | 31 | Arpa_PC12_day4_8-31-2024_Slot1-31_7704.d |
| SC_PC12_D4_3 | PC12 | Day 4 | 8/31/24 | 32 | Arpa_PC12_day4_8-31-2024_Slot1-32_7705.d |
| SC_PC12_D4_4 | PC12 | Day 4 | 8/31/24 | 33 | Arpa_PC12_day4_8-31-2024_Slot1-33_7706.d |
| SC_PC12_D4_5 | PC12 | Day 4 | 8/31/24 | 34 | Arpa_PC12_day4_8-31-2024_Slot1-34_7707.d |
| SC_PC12_D4_6 | PC12 | Day 4 | 8/31/24 | 35 | Arpa_PC12_day4_8-31-2024_Slot1-35_7708.d |
| SC_PC12_D4_7 | PC12 | Day 4 | 8/31/24 | 36 | Arpa_PC12_day4_8-31-2024_Slot1-36_7709.d |
| SC_PC12_D4_8 | PC12 | Day 4 | 8/31/24 | 37 | Arpa_PC12_day4_8-31-2024_Slot1-37_7710.d |
| SC_PC12_D4_9 | PC12 | Day 4 | 8/31/24 | 38 | Arpa_PC12_day4_8-31-2024_Slot1-38_7711.d |
| SC_PC12_D4_10 | PC12 | Day 4 | 9/1/24 | 39 | Arpa_PC12_day4_9-1-2024_Slot1-39_7712.d |
| SC_PC12_D4_11 | PC12 | Day 4 | 9/1/24 | 40 | Arpa_PC12_day4_9-1-2024_Slot1-40_7714.d |
| SC_PC12_D4_12 | PC12 | Day 4 | 9/1/24 | 41 | Arpa_PC12_day4_9-1-2024_Slot1-41_7715.d |
| SC_PC12_D4_13 | PC12 | Day 4 | 9/1/24 | 42 | Arpa_PC12_day4_9-1-2024_Slot1-42_7716.d |
| SC_PC12_D4_14 | PC12 | Day 4 | 9/1/24 | 43 | Arpa_PC12_day4_9-1-2024_Slot1-43_7717.d |
| SC_PC12_D4_15 | PC12 | Day 4 | 9/1/24 | 44 | Arpa_PC12_day4_9-1-2024_Slot1-44_7718.d |
| SC_PC12_D4_16 | PC12 | Day 4 | 9/1/24 | 45 | Arpa_PC12_day4_9-1-2024_Slot1-45_7719.d |
| SC_PC12_D4_17 | PC12 | Day 4 | 9/1/24 | 46 | Arpa_PC12_day4_9-1-2024_Slot1-46_7720.d |
| SC_PC12_D4_18 | PC12 | Day 4 | 9/1/24 | 47 | Arpa_PC12_day4_9-1-2024_Slot1-47_7721.d |
| SC_PC12_D4_19 | PC12 | Day 4 | 9/1/24 | 50 | Arpa_PC12_day4_9-1-2024_Slot1-50_7722.d |
| SC_PC12_D4_20 | PC12 | Day 4 | 9/1/24 | 51 | Arpa_PC12_day4_9-1-2024_Slot1-51_7723.d |
| SC_PC12_D4_21 | PC12 | Day 4 | 9/1/24 | 59 | Arpa_PC12_day4_9-1-2024_Slot1-59_7725.d |
| SC_PC12_D4_22 | PC12 | Day 4 | 9/1/24 | 60 | Arpa_PC12_day4_9-1-2024_Slot1-60_7726.d |
| SC_PC12_D4_23 | PC12 | Day 4 | 9/1/24 | 61 | Arpa_PC12_day4_9-1-2024_Slot1-61_7727.d |
| SC_PC12_D4_24 | PC12 | Day 4 | 9/1/24 | 62 | Arpa_PC12_day4_9-1-2024_Slot1-62_7728.d |
| SC_PC12_D4_25 | PC12 | Day 4 | 9/1/24 | 63 | Arpa_PC12_day4_9-1-2024_Slot1-63_7729.d |
| SC_PC12_D4_26 | PC12 | Day 4 | 9/1/24 | 64 | Arpa_PC12_day4_9-1-2024_Slot1-64_7730.d |
| SC_PC12_D4_27 | PC12 | Day 4 | 9/1/24 | 65 | Arpa_PC12_day4_9-1-2024_Slot1-65_7731.d |
| SC_PC12_D4_28 | PC12 | Day 4 | 9/1/24 | 66 | Arpa_PC12_day4_9-1-2024_Slot1-66_7732.d |
| SC_PC12_D4_29 | PC12 | Day 4 | 9/1/24 | 67 | Arpa_PC12_day4_9-1-2024_Slot1-67_7733.d |
| SC_PC12_D4_30 | PC12 | Day 4 | 9/1/24 | 68 | Arpa_PC12_day4_9-1-2024_Slot1-68_7734.d |
| SC_PC12_D4_31 | PC12 | Day 4 | 9/1/24 | 81 | Arpa_PC12_day4_9-1-2024_Slot1-81_7736.d |
| SC_PC12_D4_32 | PC12 | Day 4 | 9/2/24 | 82 | Arpa_PC12_day4_9-2-2024_Slot1-82_7737.d |
| SC_PC12_D4_33 | PC12 | Day 4 | 9/2/24 | 83 | Arpa_PC12_day4_9-2-2024_Slot1-83_7738.d |
| SC_PC12_D4_34 | PC12 | Day 4 | 9/2/24 | 84 | Arpa_PC12_day4_9-2-2024_Slot1-84_7739.d |
| SC_PC12_D4_35 | PC12 | Day 4 | 9/2/24 | 85 | Arpa_PC12_day4_9-2-2024_Slot1-85_7740.d |
| SC_PC12_D4_36 | PC12 | Day 4 | 9/2/24 | 86 | Arpa_PC12_day4_9-2-2024_Slot1-86_7741.d |
| SC_PC12_D4_37 | PC12 | Day 4 | 9/2/24 | 87 | Arpa_PC12_day4_9-2-2024_Slot1-87_7742.d |
| SC_PC12_D4_38 | PC12 | Day 4 | 9/2/24 | 88 | Arpa_PC12_day4_9-2-2024_Slot1-88_7746.d |
| SC_PC12_D4_39 | PC12 | Day 4 | 9/2/24 | 89 | Arpa_PC12_day4_9-2-2024_Slot1-89_7747.d |
| SC_PC12_D4_40 | PC12 | Day 4 | 9/2/24 | 90 | Arpa_PC12_day4_9-2-2024_Slot1-90_7748.d |
| SC_PC12_D4_41 | PC12 | Day 4 | 9/2/24 | 91 | Arpa_PC12_day4_9-2-2024_Slot1-91_7749.d |
| SC_PC12_D4_42 | PC12 | Day 4 | 9/2/24 | 92 | Arpa_PC12_day4_9-2-2024_Slot1-92_7750.d |
| SC_PC12_D6_01 | PC12 | Day 6 | 8/28/24 | 26 | Arpa_PC12_day6_8-28-2024_Slot1-26_7623.d |
| SC_PC12_D6_02 | PC12 | Day 6 | 8/28/24 | 27 | Arpa_PC12_day6_8-28-2024_Slot1-27_7624.d |
| SC_PC12_D6_03 | PC12 | Day 6 | 8/28/24 | 28 | Arpa_PC12_day6_8-28-2024_Slot1-28_7625.d |
| SC_PC12_D6_04 | PC12 | Day 6 | 8/28/24 | 29 | Arpa_PC12_day6_8-28-2024_Slot1-29_7626.d |
| SC_PC12_D6_05 | PC12 | Day 6 | 8/28/24 | 30 | Arpa_PC12_day6_8-28-2024_Slot1-30_7627.d |
| SC_PC12_D6_06 | PC12 | Day 6 | 8/28/24 | 31 | Arpa_PC12_day6_8-28-2024_Slot1-31_7628.d |
| SC_PC12_D6_07 | PC12 | Day 6 | 8/28/24 | 32 | Arpa_PC12_day6_8-28-2024_Slot1-32_7629.d |
| SC_PC12_D6_08 | PC12 | Day 6 | 8/28/24 | 33 | Arpa_PC12_day6_8-28-2024_Slot1-33_7630.d |
| SC_PC12_D6_09 | PC12 | Day 6 | 8/28/24 | 34 | Arpa_PC12_day6_8-28-2024_Slot1-34_7631.d |
| SC_PC12_D6_10 | PC12 | Day 6 | 8/28/24 | 35 | Arpa_PC12_day6_8-28-2024_Slot1-35_7632.d |
| SC_PC12_D6_11 | PC12 | Day 6 | 8/28/24 | 37 | Arpa_PC12_day6_8-28-2024_Slot1-37_7634.d |
| SC_PC12_D6_12 | PC12 | Day 6 | 8/28/24 | 38 | Arpa_PC12_day6_8-28-2024_Slot1-38_7635.d |
| SC_PC12_D6_13 | PC12 | Day 6 | 8/28/24 | 39 | Arpa_PC12_day6_8-28-2024_Slot1-39_7636.d |
| SC_PC12_D6_14 | PC12 | Day 6 | 8/29/24 | 40 | Arpa_PC12_day6_8-29-2024_Slot1-40_7637.d |
| SC_PC12_D6_15 | PC12 | Day 6 | 8/29/24 | 41 | Arpa_PC12_day6_8-29-2024_Slot1-41_7638.d |
| SC_PC12_D6_16 | PC12 | Day 6 | 8/29/24 | 42 | Arpa_PC12_day6_8-29-2024_Slot1-42_7639.d |
| SC_PC12_D6_17 | PC12 | Day 6 | 8/29/24 | 43 | Arpa_PC12_day6_8-29-2024_Slot1-43_7640.d |
| SC_PC12_D6_18 | PC12 | Day 6 | 8/29/24 | 44 | Arpa_PC12_day6_8-29-2024_Slot1-44_7641.d |
| SC_PC12_D6_19 | PC12 | Day 6 | 8/29/24 | 46 | Arpa_PC12_day6_8-29-2024_Slot1-46_7642.d |
| SC_PC12_D6_20 | PC12 | Day 6 | 8/29/24 | 47 | Arpa_PC12_day6_8-29-2024_Slot1-47_7643.d |
| SC_PC12_D6_21 | PC12 | Day 6 | 8/29/24 | 50 | Arpa_PC12_day6_8-29-2024_Slot1-50_7645.d |
| SC_PC12_D6_22 | PC12 | Day 6 | 8/29/24 | 51 | Arpa_PC12_day6_8-29-2024_Slot1-51_7646.d |
| SC_PC12_D6_23 | PC12 | Day 6 | 8/29/24 | 52 | Arpa_PC12_day6_8-29-2024_Slot1-52_7647.d |
| SC_PC12_D6_24 | PC12 | Day 6 | 8/29/24 | 53 | Arpa_PC12_day6_8-29-2024_Slot1-53_7648.d |
| SC_PC12_D6_25 | PC12 | Day 6 | 8/29/24 | 54 | Arpa_PC12_day6_8-29-2024_Slot1-54_7649.d |
| SC_PC12_D6_26 | PC12 | Day 6 | 8/29/24 | 55 | Arpa_PC12_day6_8-29-2024_Slot1-55_7650.d |
| SC_PC12_D6_27 | PC12 | Day 6 | 8/29/24 | 56 | Arpa_PC12_day6_8-29-2024_Slot1-56_7651.d |
| SC_PC12_D6_28 | PC12 | Day 6 | 8/29/24 | 57 | Arpa_PC12_day6_8-29-2024_Slot1-57_7652.d |
| SC_PC12_D6_29 | PC12 | Day 6 | 8/29/24 | 58 | Arpa_PC12_day6_8-29-2024_Slot1-58_7653.d |
| SC_PC12_D6_30 | PC12 | Day 6 | 8/29/24 | 59 | Arpa_PC12_day6_8-29-2024_Slot1-59_7654.d |
| SC_PC12_D6_31 | PC12 | Day 6 | 9/3/24 | 74 | Arpa_PC12_day6_9-3-2024_Slot1-74_7776.d |
| SC_PC12_D6_32 | PC12 | Day 6 | 9/3/24 | 75 | Arpa_PC12_day6_9-3-2024_Slot1-75_7777.d |
| SC_PC12_D6_33 | PC12 | Day 6 | 9/3/24 | 77 | Arpa_PC12_day6_9-3-2024_Slot1-77_7778.d |
| SC_PC12_D6_34 | PC12 | Day 6 | 9/3/24 | 79 | Arpa_PC12_day6_9-3-2024_Slot1-79_7779.d |
| SC_PC12_D6_35 | PC12 | Day 6 | 9/3/24 | 80 | Arpa_PC12_day6_9-3-2024_Slot1-80_7780.d |
| SC_PC12_D6_36 | PC12 | Day 6 | 9/3/24 | 81 | Arpa_PC12_day6_9-3-2024_Slot1-81_7781.d |
| SC_PC12_D6_37 | PC12 | Day 6 | 9/3/24 | 82 | Arpa_PC12_day6_9-3-2024_Slot1-82_7782.d |
| SC_PC12_D6_38 | PC12 | Day 6 | 9/3/24 | 83 | Arpa_PC12_day6_9-3-2024_Slot1-83_7783.d |
| SC_PC12_D6_39 | PC12 | Day 6 | 9/3/24 | 84 | Arpa_PC12_day6_9-3-2024_Slot1-84_7784.d |
| SC_PC12_D6_40 | PC12 | Day 6 | 9/3/24 | 85 | Arpa_PC12_day6_9-3-2024_Slot1-85_7785.d |
| SC_PC12_D6_41 | PC12 | Day 6 | 9/4/24 | 98 | Arpa_PC12_day6_9-4-2024_Slot1-98_7787.d |
| SC_PC12_D6_42 | PC12 | Day 6 | 9/4/24 | 99 | Arpa_PC12_day6_9-4-2024_Slot1-99_7788.d |
| SC_PC12_D6_43 | PC12 | Day 6 | 9/4/24 | 100 | Arpa_PC12_day6_9-4-2024_Slot1-100_7789.d |
| SC_PC12_D6_44 | PC12 | Day 6 | 9/4/24 | 101 | Arpa_PC12_day6_9-4-2024_Slot1-101_7790.d |
| SC_PC12_D6_45 | PC12 | Day 6 | 9/4/24 | 102 | Arpa_PC12_day6_9-4-2024_Slot1-102_7791.d |
| SC_PC12_D6_46 | PC12 | Day 6 | 9/4/24 | 103 | Arpa_PC12_day6_9-4-2024_Slot1-103_7792.d |
| SC_PC12_D6_47 | PC12 | Day 6 | 9/4/24 | 104 | Arpa_PC12_day6_9-4-2024_Slot1-104_7793.d |
| SC_PC12_D6_48 | PC12 | Day 6 | 9/4/24 | 105 | Arpa_PC12_day6_9-4-2024_Slot1-105_7794.d |
| SC_PC12_D6_49 | PC12 | Day 6 | 9/4/24 | 107 | Arpa_PC12_day6_9-4-2024_Slot1-107_7795.d |
| SC_PC12_D6_50 | PC12 | Day 6 | 9/4/24 | 108 | Arpa_PC12_day6_9-4-2024_Slot1-108_7796.d |

**Table S4.** Statistical analysis of precursor counts per single PC12 cell across differentiation stages (Day 0, Day 2, Day 4, Day 6). Each point represents an individual cell. Statistical significance was assessed using a Kruskal–Wallis test followed by Dunn’s multiple comparisons test for post hoc analysis.

|  |  |  |
| --- | --- | --- |
| Kruskal-Wallis test with Dunn's multiple comparisons test |  |  |
| Kruskal-Wallis test |  |  |
| P-value<br>(approximate<br>) | <0.0001**** |  |
| Kruskal-Wallis<br>statistic | 29.3905 |  |
| Dunn's multiple comparison test summary |  |  |
| Comparison | Adjusted P-value | Mean rank difference |
| Day 0 Undiff<br>vs Day 2<br>Diff | 0.7309ns | -13.0552 |
| Day 0 Undiff<br>vs Day 4<br>Diff | 0.002188** | 36.7167 |
| Day 0 Undiff<br>vs Day 6<br>Diff | 0.01794* | 28.7686 |

| Dunn's multiple comparison test additional details |  |  |  |  |  |  |
| --- | --- | --- | --- | --- | --- | --- |
| Comparison | Mean rank 1 | Mean rank 2 | Mean rank difference | n1 | n2 | z |
| Day 0 Undiff vs Day 2 Diff | 97.7286 | 110.7838 | -13.0552 | 35 | 37 | 1.1659 |
| Day 0 Undiff vs Day 4 Diff | 97.7286 | 61.0119 | 36.7167 | 35 | 42 | -3.3783 |
| Day 0 Undiff vs Day 6 Diff | 97.7286 | 68.96 | 28.7686 | 35 | 50 | -2.7489 |
| Descriptive statistics |  |  |  |  |  |  |
| Category | Mean | SD | SEM |  |  |  |
| Day 0 Undiff | 9152.8571 | 3964.2068 | 670.0733 |  |  |  |
| Day 2 Diff | 12483.8649 | 8441.0081 | 1387.6932 |  |  |  |
| Day 4 Diff | 6205.3571 | 3342.2484 | 515.7201 |  |  |  |
| Day 6 Diff | 6644.86 | 2676.873 | 378.567 |  |  |  |
| Category | Minimum | 25th percentile | Median | 75th percentile | Maximum |  |
| Day 0 Undiff | 1054 | 6449 | 8542 | 12805 | 17630 |  |
| Day 2 Diff | 0 | 6701 | 10923 | 14320.5 | 35677 |  |
| Day 4 Diff | 837 | 3404.75 | 5652.5 | 7915.25 | 15347 |  |
| Day 6 Diff | 2444 | 4279.75 | 6328 | 8644 | 12050 |  |
| Shapiro-Wilk normality test |  |  |  |  |  |  |
| Category | W | p-value | Passed<br>normality test<br>(alpha=0.05)<br>? | n |  |  |
| Day 0 Undiff | 0.9762 | 0.6329ns | Yes | 35 |  |  |
| Day 2 Diff | 0.8753 | 0.0006489**<br>* | No | 37 |  |  |
| Day 4 Diff | 0.9264 | 0.009834** | No | 42 |  |  |
| Day 6 Diff | 0.9571 | 0.07295ns | Yes | 50 |  |  |
| Levene test of equal variances |  |  |  |  |  |  |
| F | DFn | DFd | p-value | Are the SDs significantly different? |  |  |
| 8.2603 | 3 | 160 | <0.0001*** | Yes |  |  |

**Table S5.**
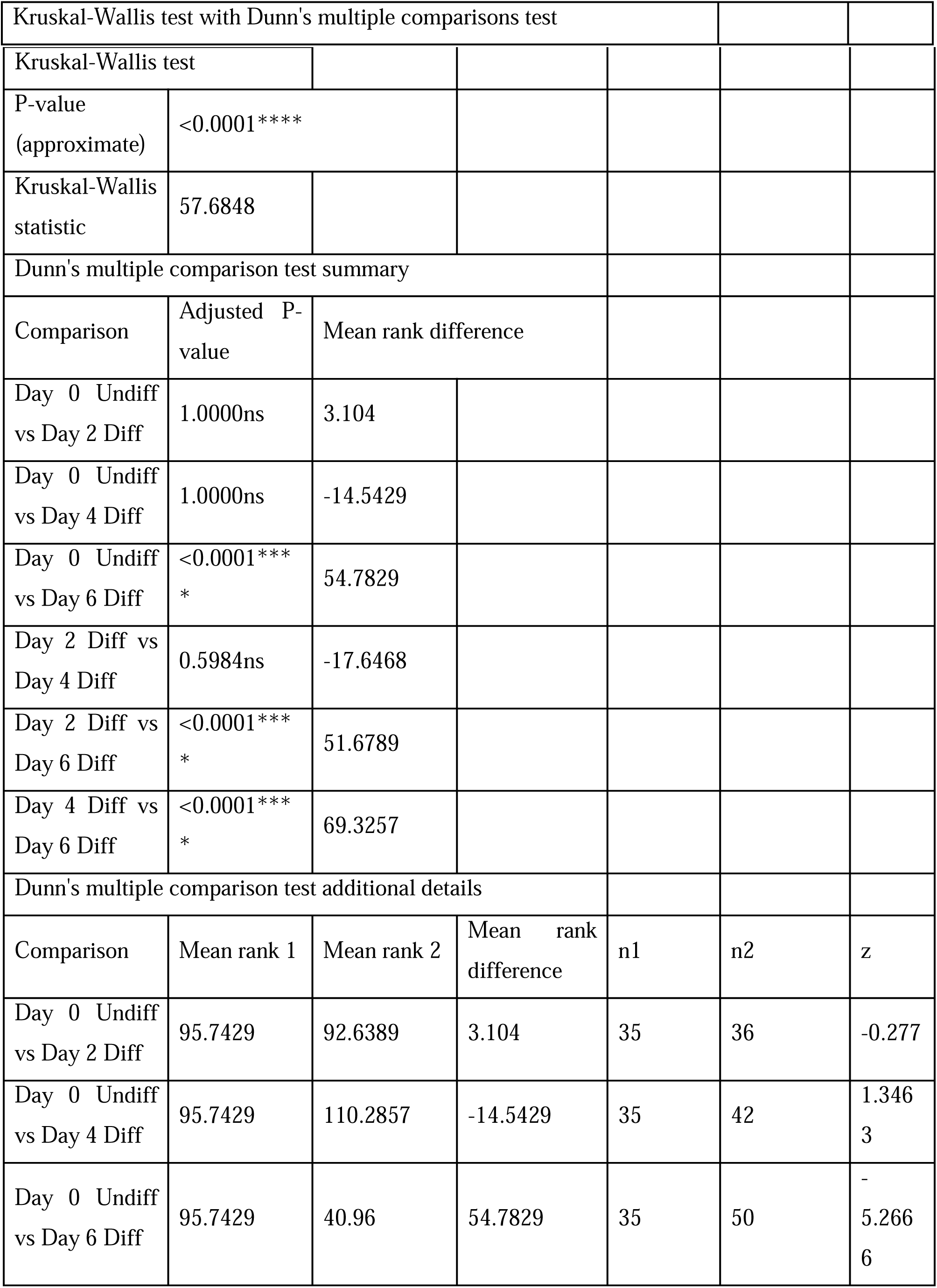
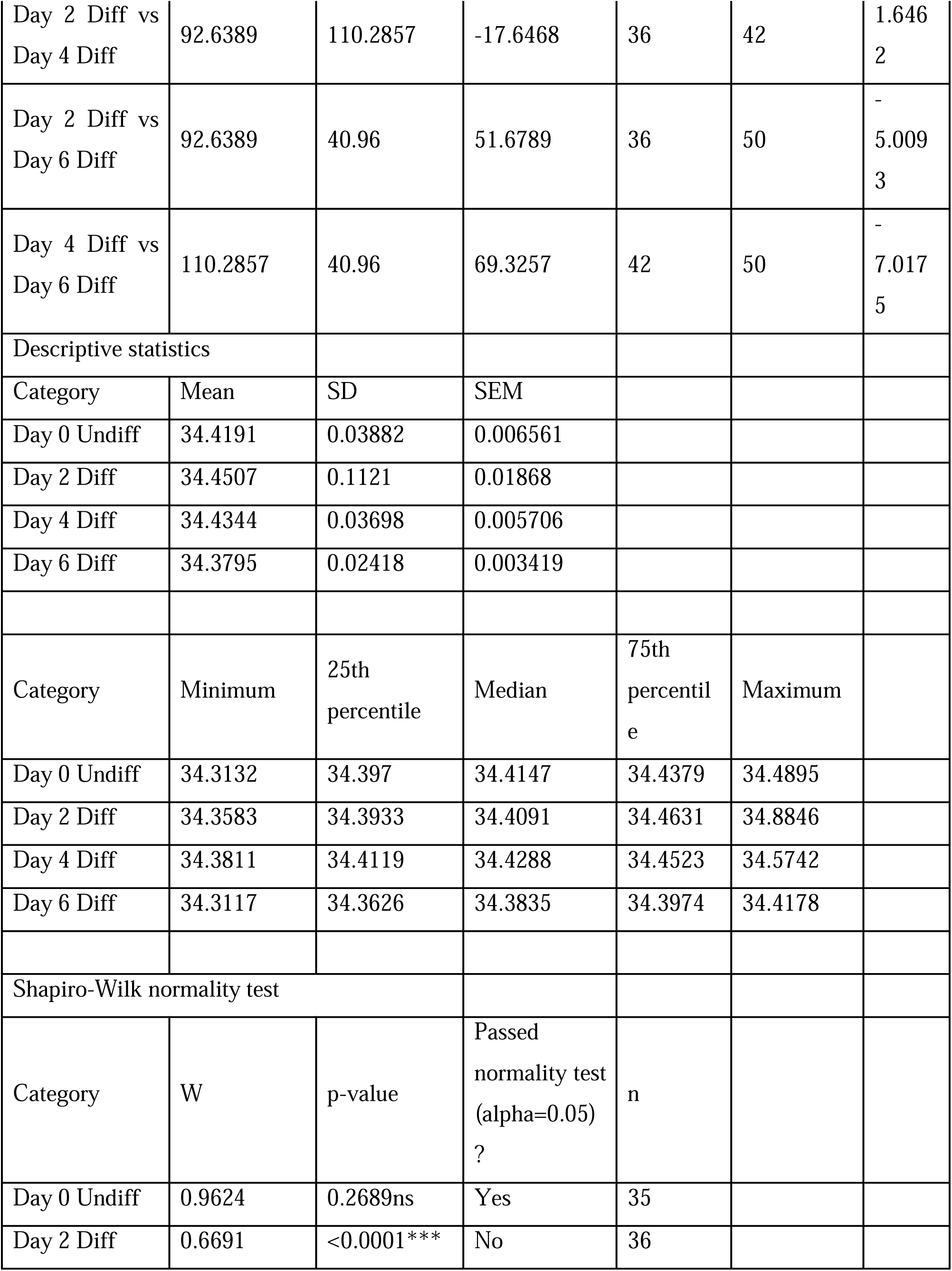
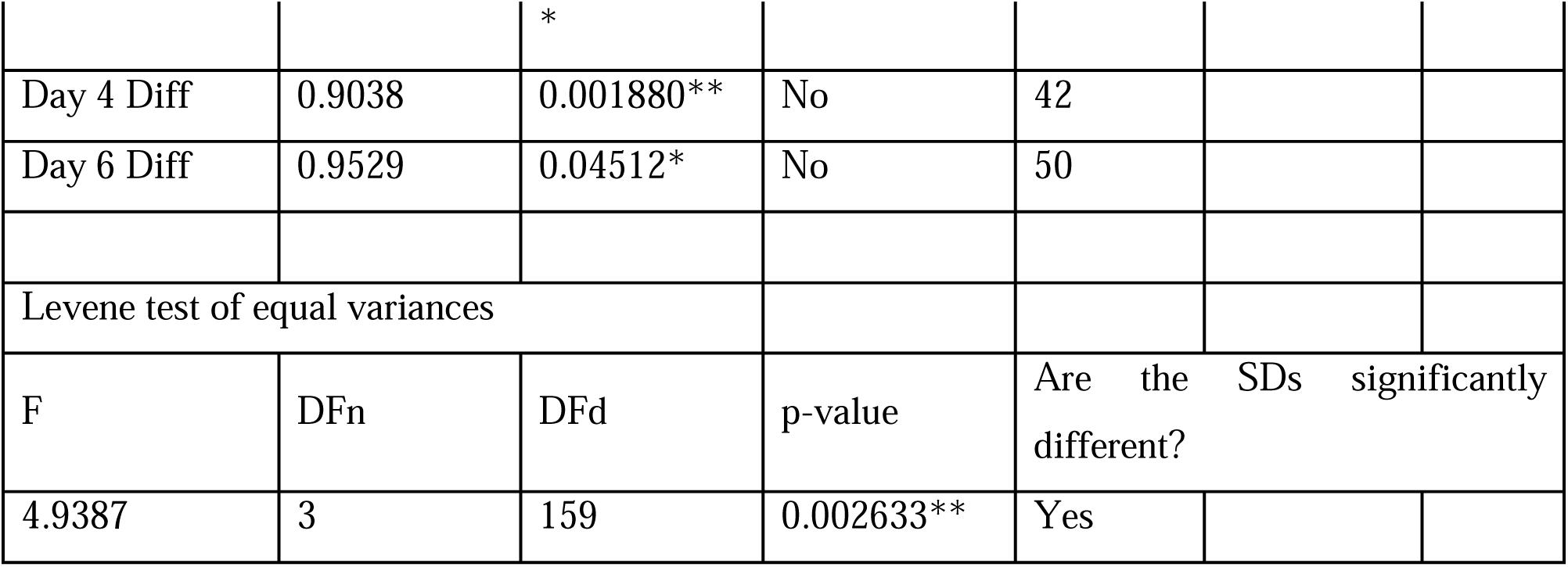
Statistical analysis of log intensity distributions of precursors across differentiation stages.

**Table S6.** Top 50 proteins ranked high in bulk but low in single-cell analyses. Proteins are ordered by the difference between bulk and single-cell percentile scores (Δ Bulk − Single-cell percentile). Higher percentiles indicate greater relative abundance within each dataset. These proteins show elevated abundance in bulk compared to single-cell measurements.

| Protein | Bulk_ | SingleCell_ | Bulk_ | SingleCell_ | Delta_ |
| --- | --- | --- | --- | --- | --- |

|  | **Rank** | **Rank** | **Percentile** | **Percentile** | **(Bulk_**<br>**minus_SC)** |
| --- | --- | --- | --- | --- | --- |
| Ccdc9 | 28 | 3707 | 0.9973 | 0.2414 | 0.7559 |
| Sacs | 259 | 4763 | 0.9741 | 0.0252 | 0.9489 |
| Rps7-ps23 | 266 | 4057 | 0.9734 | 0.1697 | 0.8037 |
| Jakmip1 | 283 | 4228 | 0.9717 | 0.1347 | 0.8370 |
| Ppp2r1b | 373 | 4166 | 0.9626 | 0.1474 | 0.8152 |
| RGD1563551 | 377 | 4244 | 0.9622 | 0.1314 | 0.8308 |
| Rpl29 | 379 | 4558 | 0.9620 | 0.0671 | 0.8949 |
| Ndufab1 | 383 | 4407 | 0.9616 | 0.0981 | 0.8636 |
| ENSRNOG00000069381 | 454 | 4381 | 0.9545 | 0.1034 | 0.8511 |
| Cpne7 | 754 | 4174 | 0.9243 | 0.1458 | 0.7786 |
| Pacsin3 | 811 | 4395 | 0.9186 | 0.1005 | 0.8181 |
| Dnajc5 | 831 | 4465 | 0.9166 | 0.0862 | 0.8304 |
| Ppa2 | 934 | 4799 | 0.9063 | 0.0178 | 0.8884 |
| Rpl37-ps4 | 1041 | 4703 | 0.8955 | 0.0375 | 0.8580 |
| Tmem120a | 1044 | 4377 | 0.8952 | 0.1042 | 0.7910 |
| Trappc8 | 1147 | 4072 | 0.8849 | 0.1666 | 0.7182 |
| Jpt1 | 1216 | 4121 | 0.8779 | 0.1566 | 0.7213 |
| Zc3h4 | 1267 | 4214 | 0.8728 | 0.1376 | 0.7352 |
| Smarce1 | 1343 | 4482 | 0.8652 | 0.0827 | 0.7825 |
| Mbd3 | 1420 | 4187 | 0.8574 | 0.1431 | 0.7143 |
| Klhl11 | 1483 | 4326 | 0.8511 | 0.1146 | 0.7365 |
| Pgs1 | 1542 | 4233 | 0.8452 | 0.1337 | 0.7115 |
| Cog4 | 1570 | 4343 | 0.8424 | 0.1112 | 0.7312 |
| Prnp | 1602 | 4348 | 0.8391 | 0.1101 | 0.7290 |
| Adi1 | 1609 | 4336 | 0.8384 | 0.1126 | 0.7259 |
| Slirp | 1630 | 4811 | 0.8363 | 0.0154 | 0.8210 |
| Slc35b2 | 1679 | 4435 | 0.8314 | 0.0923 | 0.7391 |
| Gcc1 | 1755 | 4792 | 0.8238 | 0.0192 | 0.8045 |
| RGD1560212 | 1791 | 4342 | 0.8202 | 0.1114 | 0.7088 |
| Cend1 | 1839 | 4362 | 0.8153 | 0.1073 | 0.7081 |
| Ptcd3 | 1886 | 4402 | 0.8106 | 0.0991 | 0.7115 |
| Ttc9c | 1934 | 4739 | 0.8058 | 0.0301 | 0.7757 |
| Erg28 | 1944 | 4458 | 0.8048 | 0.0876 | 0.7172 |
| Pex16 | 1962 | 4819 | 0.8030 | 0.0137 | 0.7893 |
| Mbnl3 | 2009 | 4529 | 0.7983 | 0.0731 | 0.7252 |
| Tmem245 | 2064 | 4547 | 0.7927 | 0.0694 | 0.7233 |
| Mapk8ip3 | 2083 | 4489 | 0.7908 | 0.0813 | 0.7095 |
| Elp2 | 2088 | 4698 | 0.7903 | 0.0385 | 0.7518 |
| Trim67 | 2158 | 4511 | 0.7833 | 0.0768 | 0.7065 |
| Mrpl3 | 2180 | 4756 | 0.7811 | 0.0266 | 0.7545 |
| Baiap2 | 2181 | 4741 | 0.7810 | 0.0297 | 0.7513 |
| Gk | 2223 | 4514 | 0.7768 | 0.0762 | 0.7006 |
| Snx30 | 2362 | 4843 | 0.7628 | 0.0088 | 0.7540 |
| Pick1 | 2423 | 4604 | 0.7567 | 0.0577 | 0.6989 |
| R3hdm1 | 2424 | 4673 | 0.7566 | 0.0436 | 0.7130 |
| Washc2 | 2472 | 4821 | 0.7517 | 0.0133 | 0.7384 |
| Ppp2r5b | 2487 | 4869 | 0.7502 | 0.0035 | 0.7467 |
| Cwc25 | 2488 | 4610 | 0.7501 | 0.0565 | 0.6936 |
| ENSRNOG00000062602 | 2536 | 4788 | 0.7453 | 0.0201 | 0.7252 |
| Sugp1 | 2650 | 4772 | 0.7338 | 0.0233 | 0.7105 |

**Table S7.**
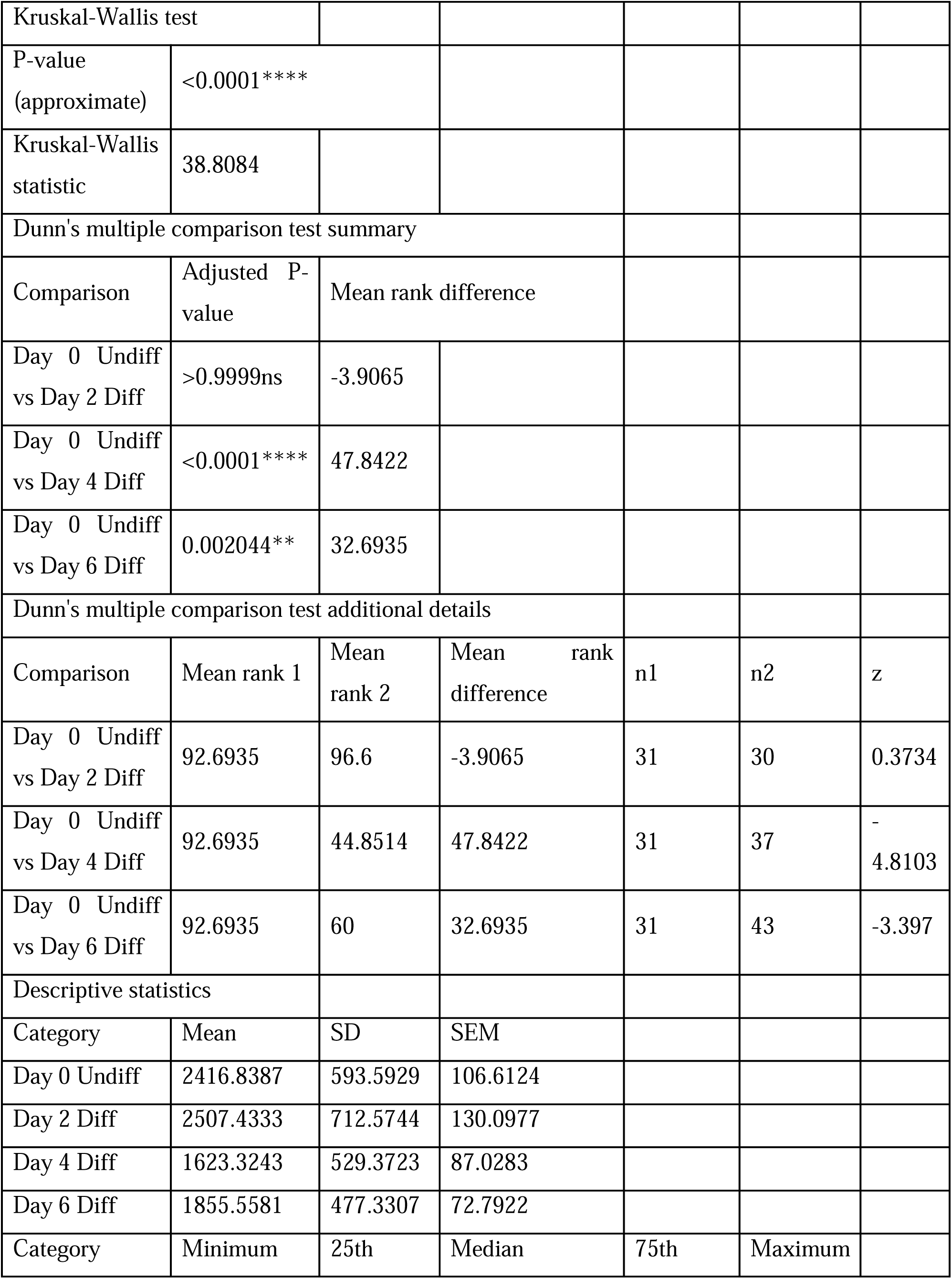
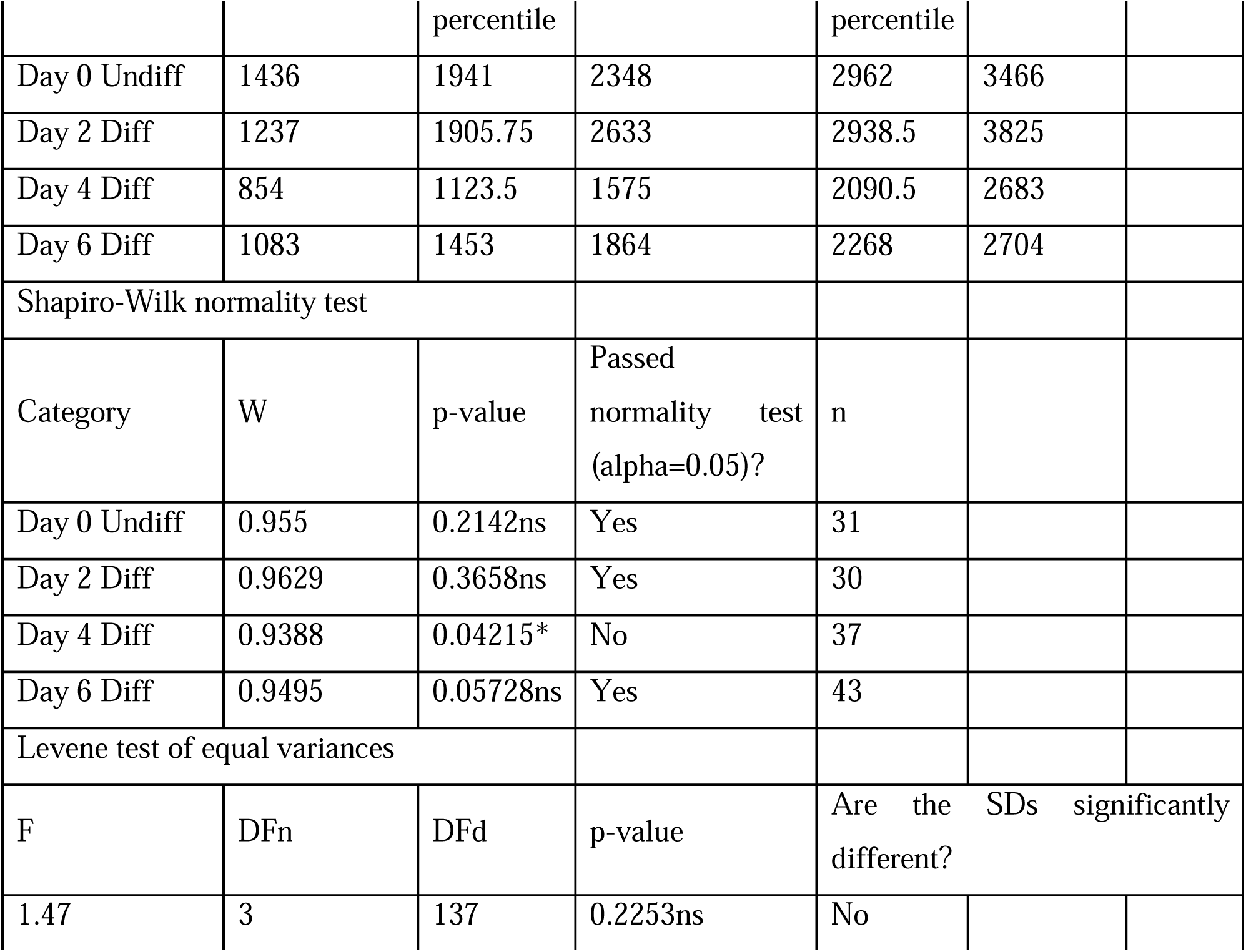
Comparison of protein abundance distributions across differentiation stages. Protein abundance distributions for Day 0, Day 2, Day 4, and Day 6 single-cell samples were compared using a Kruskal–Wallis nonparametric test. A significant difference was observed among the groups (Kruskal–Wallis H = 38.808, p < 0.001), indicating stage-dependent shifts in protein abundance distributions.

|  |  |  |  |  |  |  |
| --- | --- | --- | --- | --- | --- | --- |
| Kruskal-Wallis test |  |  |  |  |  |  |
| P-value<br>(approximate) | <0.0001**** |  |  |  |  |  |
| Kruskal-Wallis<br>statistic | 38.8084 |  |  |  |  |  |
| Dunn's multiple comparison test summary |  |  |  |  |  |  |
| Comparison | Adjusted P-value | Mean rank difference |  |  |  |  |
| Day 0 Undiff<br>vs Day 2 Diff | >0.9999ns | -3.9065 |  |  |  |  |
| Day 0 Undiff<br>vs Day 4 Diff | <0.0001**** | 47.8422 |  |  |  |  |
| Day 0 Undiff<br>vs Day 6 Diff | 0.002044** | 32.6935 |  |  |  |  |
| Dunn's multiple comparison test additional details |  |  |  |  |  |  |
| Comparison | Mean rank 1 | Mean rank 2 | Mean rank difference | n1 | n2 | z |
| Day 0 Undiff<br>vs Day 2 Diff | 92.6935 | 96.6 | -3.9065 | 31 | 30 | 0.3734 |
| Day 0 Undiff<br>vs Day 4 Diff | 92.6935 | 44.8514 | 47.8422 | 31 | 37 | -4.8103 |
| Day 0 Undiff<br>vs Day 6 Diff | 92.6935 | 60 | 32.6935 | 31 | 43 | -3.397 |
| Descriptive statistics |  |  |  |  |  |  |
| Category | Mean | SD | SEM |  |  |  |
| Day 0 Undiff | 2416.8387 | 593.5929 | 106.6124 |  |  |  |
| Day 2 Diff | 2507.4333 | 712.5744 | 130.0977 |  |  |  |
| Day 4 Diff | 1623.3243 | 529.3723 | 87.0283 |  |  |  |
| Day 6 Diff | 1855.5581 | 477.3307 | 72.7922 |  |  |  |
| Category | Minimum | 25th | Median | 75th | Maximum |  |

|  |  |  |  |  |  |
| --- | --- | --- | --- | --- | --- |
|  |  | percentile |  | percentile |  |
| Day 0 Undiff | 1436 | 1941 | 2348 | 2962 | 3466 |
| Day 2 Diff | 1237 | 1905.75 | 2633 | 2938.5 | 3825 |
| Day 4 Diff | 854 | 1123.5 | 1575 | 2090.5 | 2683 |
| Day 6 Diff | 1083 | 1453 | 1864 | 2268 | 2704 |
| Shapiro-Wilk normality test |  |  |  |  |  |
| Category | W | p-value | Passed normality test (alpha=0.05)? | n |  |
| Day 0 Undiff | 0.955 | 0.2142ns | Yes | 31 |  |
| Day 2 Diff | 0.9629 | 0.3658ns | Yes | 30 |  |
| Day 4 Diff | 0.9388 | 0.04215* | No | 37 |  |
| Day 6 Diff | 0.9495 | 0.05728ns | Yes | 43 |  |
| Levene test of equal variances |  |  |  |  |  |
| F | DFn | DFd | p-value | Are the SDs significantly different? |  |
| 1.47 | 3 | 137 | 0.2253ns | No |  |

**Table S8.**
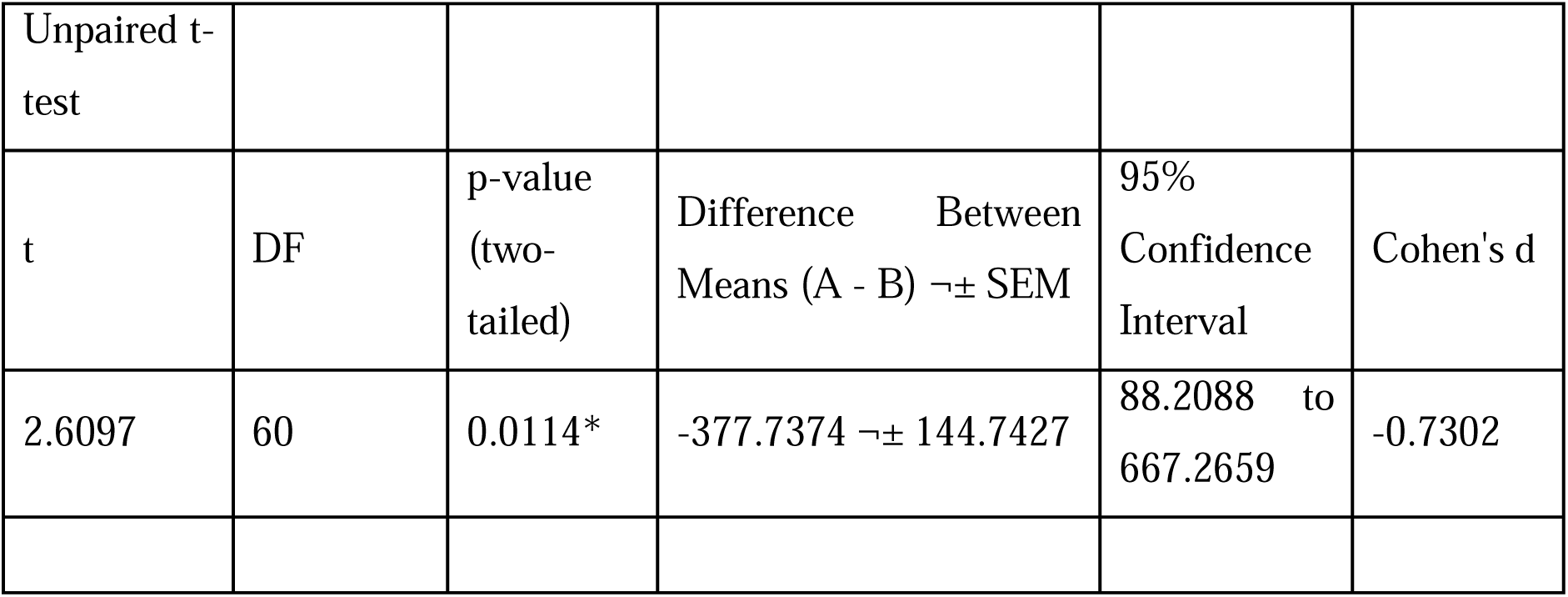
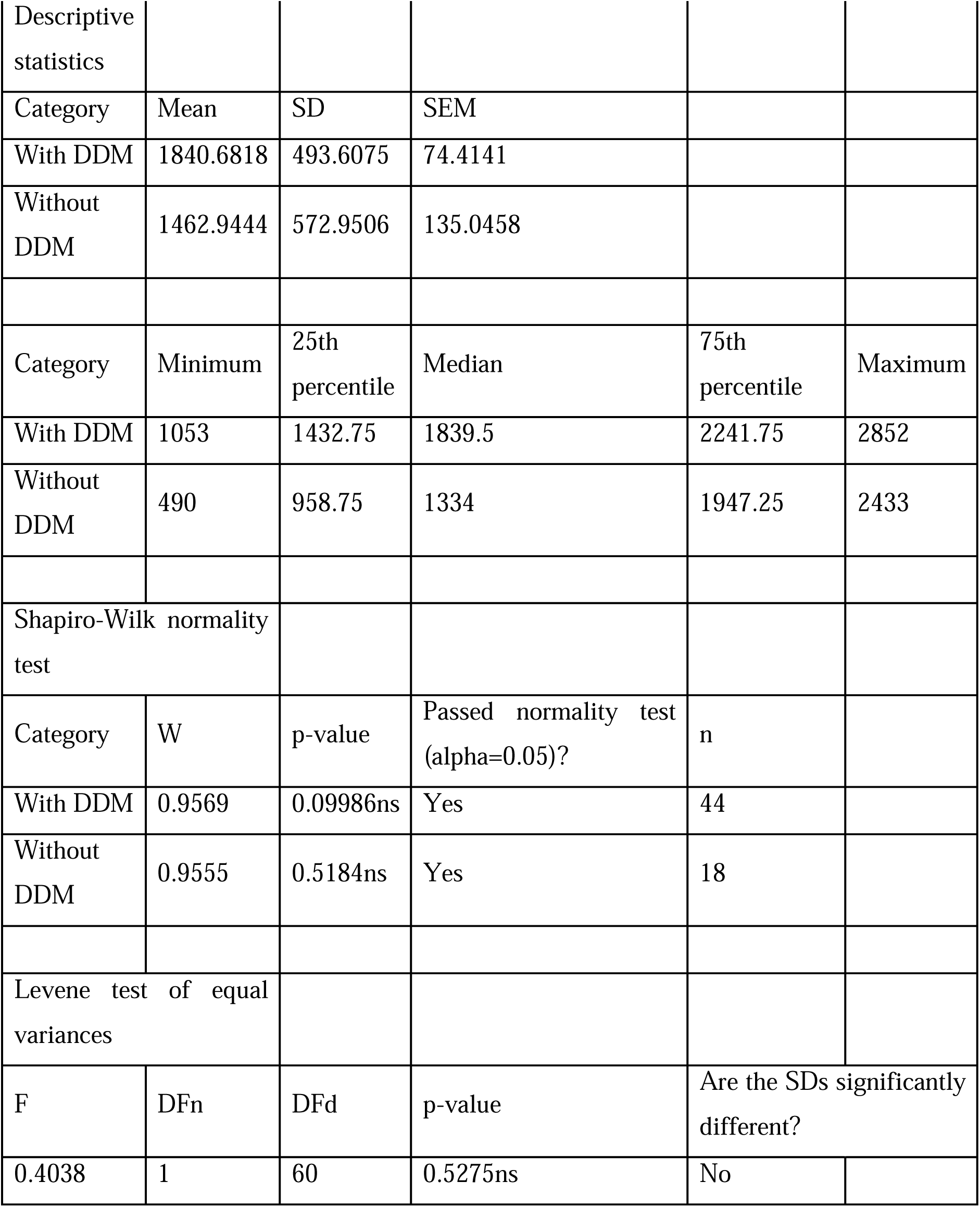
Statistical analysis of protein abundance with and without DDM. Unpaired two-tailed *t*-test (α = 0.05) was used following Shapiro–Wilk normality and Levene’s variance tests; effect size reported as Cohen’s *d*. *p* < 0.05 was considered significant.

**Table S9.**
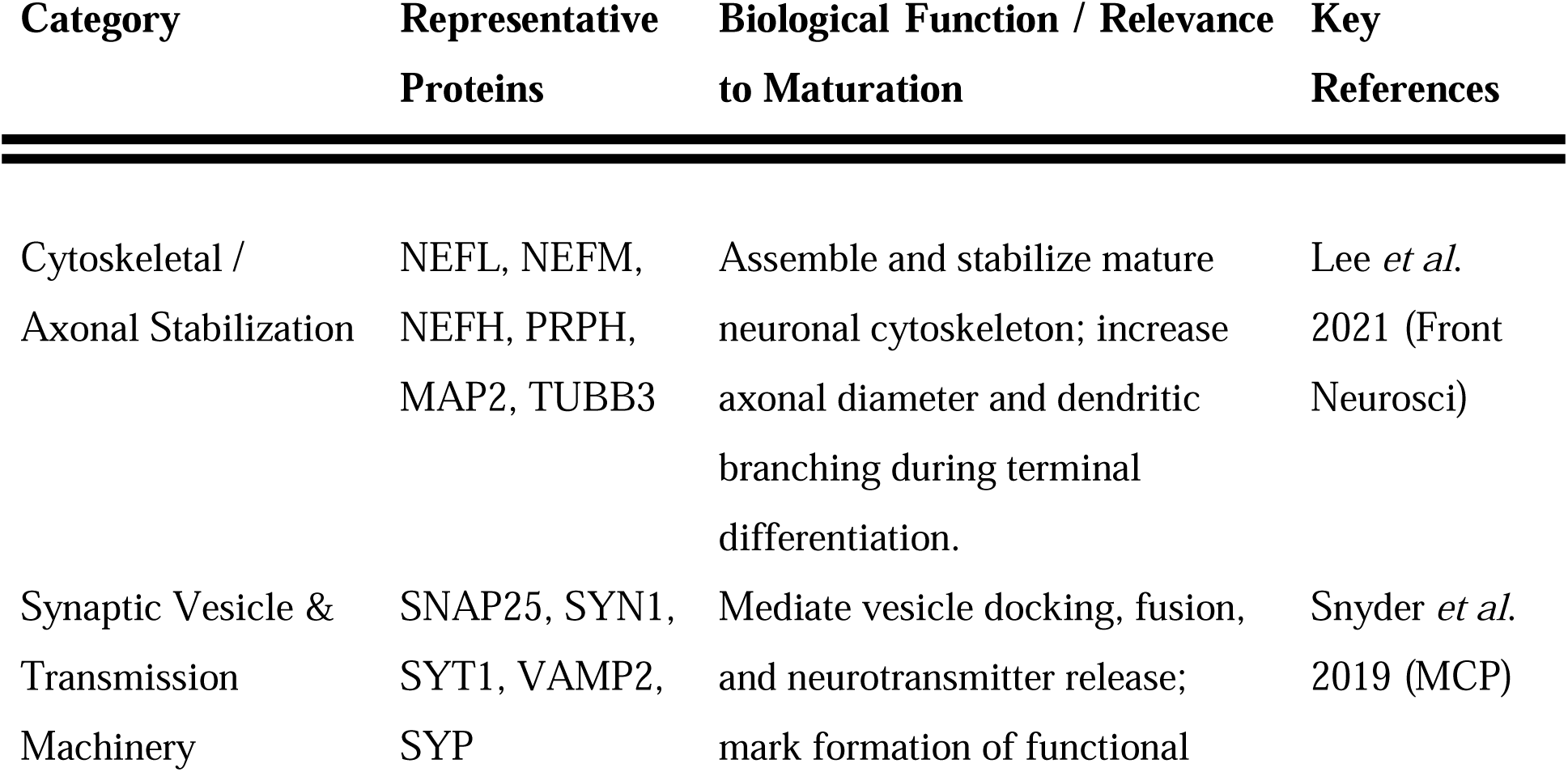
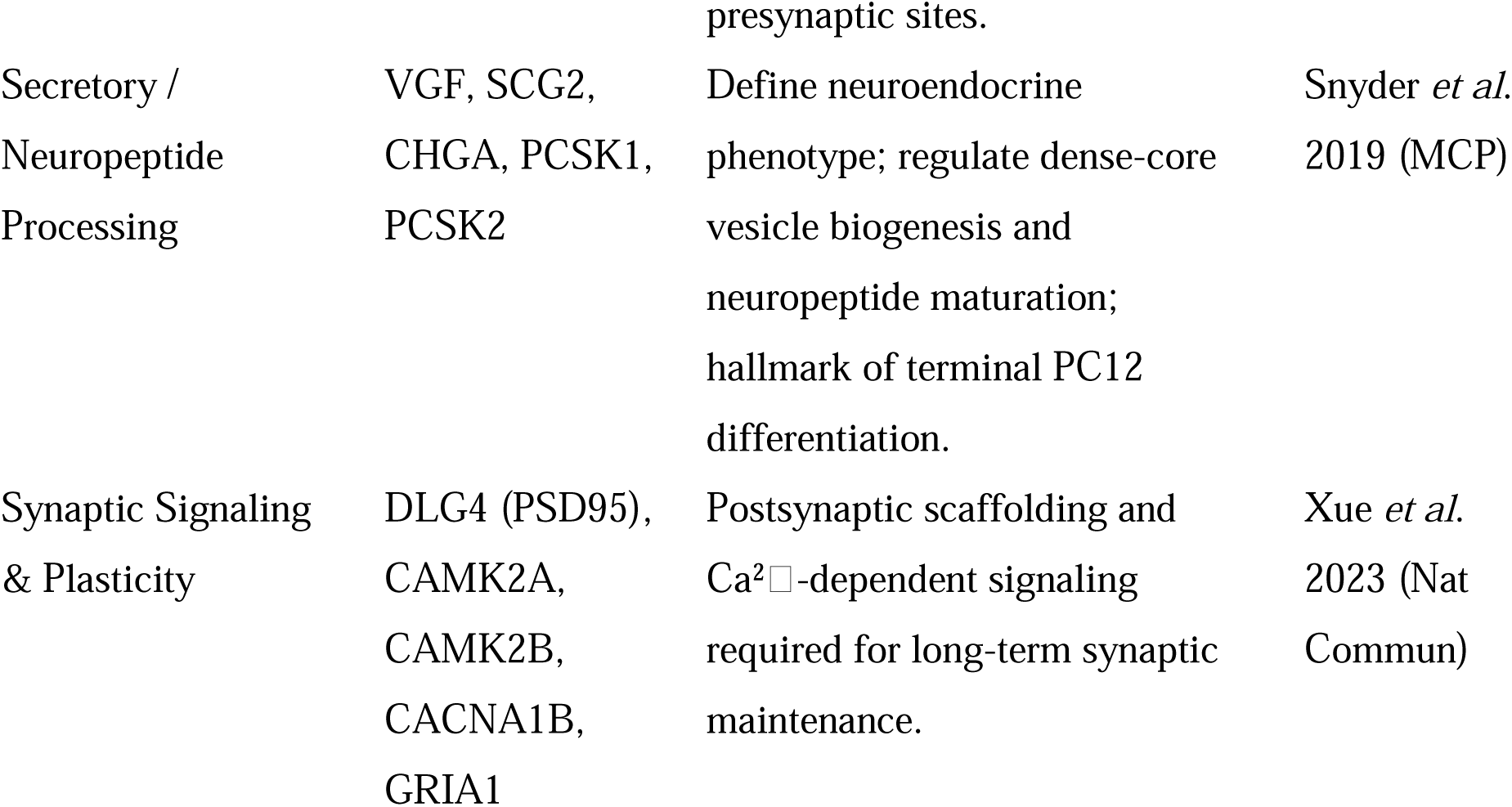
Neuronal maturation markers detected in late PC12 differentiation (Day 4–6).

**Table S10.** Proteins contributing to Day 2 subgroup separation identified by PCA. Proteins with significant z-scores and p-values along PC1 were identified from PCA of Day 2 single-cell proteomic data. PC1 contribution indicates the direction and magnitude of association with the two Day 2 subgroups. Functional annotations highlight enrichment in translation, RNA regulation, and intracellular trafficking pathways.

| Gene | PC1<br>Contribution | Z-score | p-value | Interpretation (PC1<br>Direction) | Primary Functional<br>Category |
| --- | --- | --- | --- | --- | --- |
| Nat10 | 0.0584 | 2.02 | 0.0436 | Enriched in PC1-positive subgroup | Translation / RNA<br>regulation |
| Eprs1 | −0.0527 | −1.98 | 0.0472 | Enriched in PC1-negative subgroup | Translation / tRNA<br>charging |
| Eif3f | −0.0559 | −2.10 | 0.0359 | Enriched in PC1-negative subgroup | Translation initiation |
| Arl1 | −0.0551 | −2.07 | 0.0384 | Enriched in PC1-negative subgroup | Vesicle trafficking |

